# Fatty Acid Synthase associates with nuclear-derived cytoplasmic dsRNA molecules and influences antiviral innate immune response

**DOI:** 10.1101/2025.07.16.662511

**Authors:** Charline Pasquier, Mélanie Messmer, Lise Moroge, Lisanne Knol, Johana Chicher, Richard Patryk Ngondo, Sébastien Pfeffer, Erika Girardi

**Affiliations:** UPR 9002 – Architecture et Réactivité de l’ARN, CNRS, Institut de Biologie Moléculaire et Cellulaire, 2 allée Konrad Roentgen, 67084 Strasbourg, France; Plateforme Protéomique Strasbourg-Esplanade, CNRS UAR1589, Institut de Biologie Moléculaire et Cellulaire, 2 allée Konrad Roentgen, 67084 Strasbourg, France

## Abstract

Cytoplasmic double-stranded RNA (dsRNA) is a hallmark of viral infection. It triggers innate immune responses after recognition by pathogen recognition receptors (PRRs) and leads to the production of interferons. DsRNAs arising from endogenous transcripts can also contribute to immune regulation, depending on their abundance, localization and processing. However, the mechanisms of their immunogenic potential and interactions with host proteins remain poorly understood.

Fatty Acid Synthase (FASN) is a key metabolic enzyme involved in *de novo* lipid synthesis, implicated in the regulation of cellular growth and increasingly recognized for its pro-viral functions. Here, we investigate the impact of FASN on endogenous dsRNA dynamics in human cells and its potential role in innate immune sensing. We show that FASN depletion increases the cytoplasmic accumulation of endogenous dsRNAs and promotes their enrichment in proximity to mitochondria in HCT116 cells. Transcriptomic analyses reveal that FASN deficiency is not only associated with alterations in metabolic pathways, but also with increased expression of inflammation-related genes, such as the interferon-stimulated gene (ISG) IFIT1. Combining dsRNA pulldowns with mass spectrometry and RNA-seq demonstrates that FASN associates with a subset of cytoplasmic dsRNAs derived from nuclear-encoded transcripts under basal conditions and that its association increases when endo-dsRNA levels are experimentally elevated. FASN-deficient cells display enhanced responsiveness to exogenous dsRNA stimulation with both poly I:C and viral RNA. In addition, FASN depletion restricts replication of Sindbis virus and is associated with sustained dsRNA accumulation despite reduced viral RNA levels, supporting a link between FASN activity, dsRNA regulation and antiviral responses. Our findings identify FASN as a critical regulator of endogenous dsRNA accumulation and sensing, revealing a potential role in innate immune response modulation.

## Introduction

Endogenous double-stranded RNAs (endo-dsRNAs) have emerged as critical modulators of innate immunity, with significant implications for the pathogenesis of autoimmune disorders^1^. These molecules primarily derive from nuclear retroelements, including SINEs, as well as from mitochondrial transcripts generated through bidirectional transcription of the mitochondrial genome ^1,2^. Under physiological conditions, the accumulation of dsRNAs in the cytoplasm is tightly controlled. However, when these regulatory constraints are breached, cytoplasmic endo-dsRNAs are recognized by PRRs ^1^such as RIG-I-like receptors (RLRs) in a sequence-independent and structure-dependent manner ^3^. Engagement of RLRs initiates a biphasic interferon (IFN) response: an initial phase involving the cytoplasmic sensing of dsRNA and subsequent secretion of IFNs, followed by a second phase wherein secreted IFNs bind their cognate receptors and induce the transcription of interferon-stimulated genes (ISGs)^4^. Type I and type III IFNs are robustly induced upon detection of viral nucleic acids in most human cell types, although the type III IFN response is more specific to epithelial cells ^5^. Type I IFNs include 17 members in humans, compared to the 4 type III IFN members, namely IFNL1, IFNL2, IFNL3, and IFNL4. While type I IFNs trigger a stronger and faster response that resolves quickly, type III IFNs induce a weaker response that persists longer. Despite these differences and their receptor specificity, both pathways activate overlapping sets of ISGs ^4^, such as ISG15 ^6^, IFIT1 ^7^ or OAS3 ^8^. Given the deleterious consequences of cytoplasmic dsRNA accumulation, multiple and complementary regulatory mechanisms have evolved to control aberrant accumulation of cytoplasmic endo-dsRNAs and preserve immune homeostasis ^1^. For instance, nuclear genes containing repeated elements as a source of endo-dsRNAs are subject to stringent epigenetic repression ^9,10^. An additional layer of post-transcriptional safeguard against immunogenic dsRNA expression relies on RNA-binding proteins (RBPs)^1^. Among these, hnRNPC favors excision of SINE-enriched introns, thereby preventing the export of dsRNA-containing transcripts ^11^. PNPT1 restricts the cytoplasmic leakage of mitochondrial dsRNAs ^12^. ADAR1 modulates dsRNA immunogenicity through adenosine-to-inosine (A-to-I) editing, which disrupt double-stranded structure and reduces immune recognition ^13^.

Microbial infections can perturb the tightly regulated endo-dsRNA homeostasis, promoting enhancement of innate immune signaling. Lipopolysaccharide (LPS) stimulation, which mimics bacterial infection, increases A-to-I editing, possibly reflecting changes in the dsRNA landscape ^14^. In agreement, LPS treatment causes mitochondrial dsRNA release into the cytoplasm, thereby activating the IFN signaling cascade ^15^. Moreover, influenza A virus (IAV) infection induces derepression of nuclear retroelements, particularly inverted Alu repeats (IRAlus) within the 3’ UTR of host transcripts, generating immunostimolatory endo-dsRNAs able to potentiate the antiviral response. Of note, the viral protein NS1 sequesters those endo-dsRNAs to limit their immunogenicity and therefore counteracts antiviral defense^16^.

These observations indicate that RBPs represent an important class of regulators of dsRNA homeostasis. In order to identify host RBPs able to control endo-dsRNA accumulation and immunogenicity upon infection, we previously applied a dsRNA capture approach coupled with mass spectrometry (DRIMS) during Sindbis virus (SINV) infection ^17^. Within the dsRNA interactome, the human Fatty Acid Synthase (FASN) was retrieved in both infected and uninfected cells, suggesting that it could associate with dsRNA of endogenous and viral origin.

FASN is a 270-kDa multifunctional cytoplasmic enzyme responsible for the *de novo* synthesis of palmitate from acetyl-CoA and malonyl-CoA in the presence of NADH ^18^, a key precursor for numerous fatty acids and triglycerides. FASN is composed of two identical subunits, each containing seven catalytic domains: β-ketoacyl synthase (KS), malonyl/acetyltransferase (MAT), dehydrogenase (DH), enoyl reductase (ER), β -ketoacyl reductase (KR), acyl carrier protein (ACP) and thioesterase (TE) ^19^.

Beyond its central role in lipid metabolism, FASN supports membrane plasticity, energy storage and cellular proliferation ^20^ . Consistent with these functions, FASN is frequently upregulated in various cancers ^21,22^, where it promotes tumor growth and survival by enhanced lipid synthesis ^23^. FASN has also been reported to modulate immune responses, as its overexpression is associated with the downregulation of immune-related gene expression at the mRNA level ^24^. The palmitate produced by FASN also serves as a substrate for protein palmitoylation, a post-translational modification essential for the proper localization and function of many membrane-associated proteins ^25^. These metabolic activities can also support viral replication, as demonstrated for several viruses ^26–28^, including Chikungunya virus (CHIKV), where palmitoylation of the viral protein Nsp1 is critical for efficient replication ^29^. While FASN binding to CHIKV viral RNA has been demonstrated ^30^, it has not yet been shown whether it can associate with endo-dsRNA or modulate dsRNA-mediated innate immune signaling.

In this study, we observed a cytoplasmic accumulation of endogenous dsRNAs in proximity to mitochondria in FASN knock-out (KO) HCT116 cells. Transcriptomic analysis shows alterations in metabolic pathways and increased expression of inflammatory genes, as well as an increased type III IFN production in FASN-depleted cells. Using dsRNA-IP and FASN-RNA immunoprecipitation (RIP) approaches, we analyze the dsRNA transcriptome in FASN KO and control conditions and we further identify the subset of FASN-associated RNAs. Comparative dsRNA profiling reveals minor differences in dsRNA composition between FASN KO and control cells, supporting the hypothesis that FASN primarily influences exposure of a subset of dsRNAs, rather than global dsRNA production. We also demonstrate that FASN–dsRNA association increases under conditions of induced endo-dsRNA accumulation, such as 5-AZA treatment in ADAR1 KO HCT116 cells, by proteomics and proximity ligation assay (PLA). Finally, our results show that FASN KO cells exhibit heightened sensitivity to exogenous dsRNA stimulation, with increased IFN production and stronger induction of ISGs upon poly I:C treatment. In line with that, FASN-deficient cells possess an enhanced ISG induction upon SINV infection and restrict viral replication. Together, these results identify FASN as a novel regulator of endo-dsRNA accessibility and innate immune responsiveness, linking a metabolic enzyme to the control of RNA sensing pathways without directly altering dsRNA abundance.

## Results

### FASN depletion leads to an accumulation of cytoplasmic endo-dsRNAs

To investigate whether FASN impacts immune gene expression and endo-dsRNA biology, we generated FASN KO in human colorectal cancer HCT116 cells using the CRISPR/Cas9-mediated genome editing (**Figure 1A**). HCT116 cells stably expressing *Streptococcus pyogenes* (*Sp*)Cas9 were transduced with lentiviral vectors encoding either a non-targeting control guide (cr)RNA, or a sgRNA targeting FASN exon 7. Following selection, FASN KO cells were isolated and compared to control cells expressing the non-targeting sgRNA (CTRL HCT116 cells). The KO of FASN was confirmed to be due to a 2-nt addition at the site of cleavage (**Figure 1A**). Loss of FASN protein expression was confirmed by western blot (**Figure 1B**). We next assessed the impact of FASN depletion on cell proliferation by MTT assay over 72 hours and measured a significant reduction in cell growth compared to CTRL HCT116 cells (**Figure S1A**), consistent with previous reports showing that FASN inhibition impairs cell growth ^23,24,31,32^.

**Figure 1.**
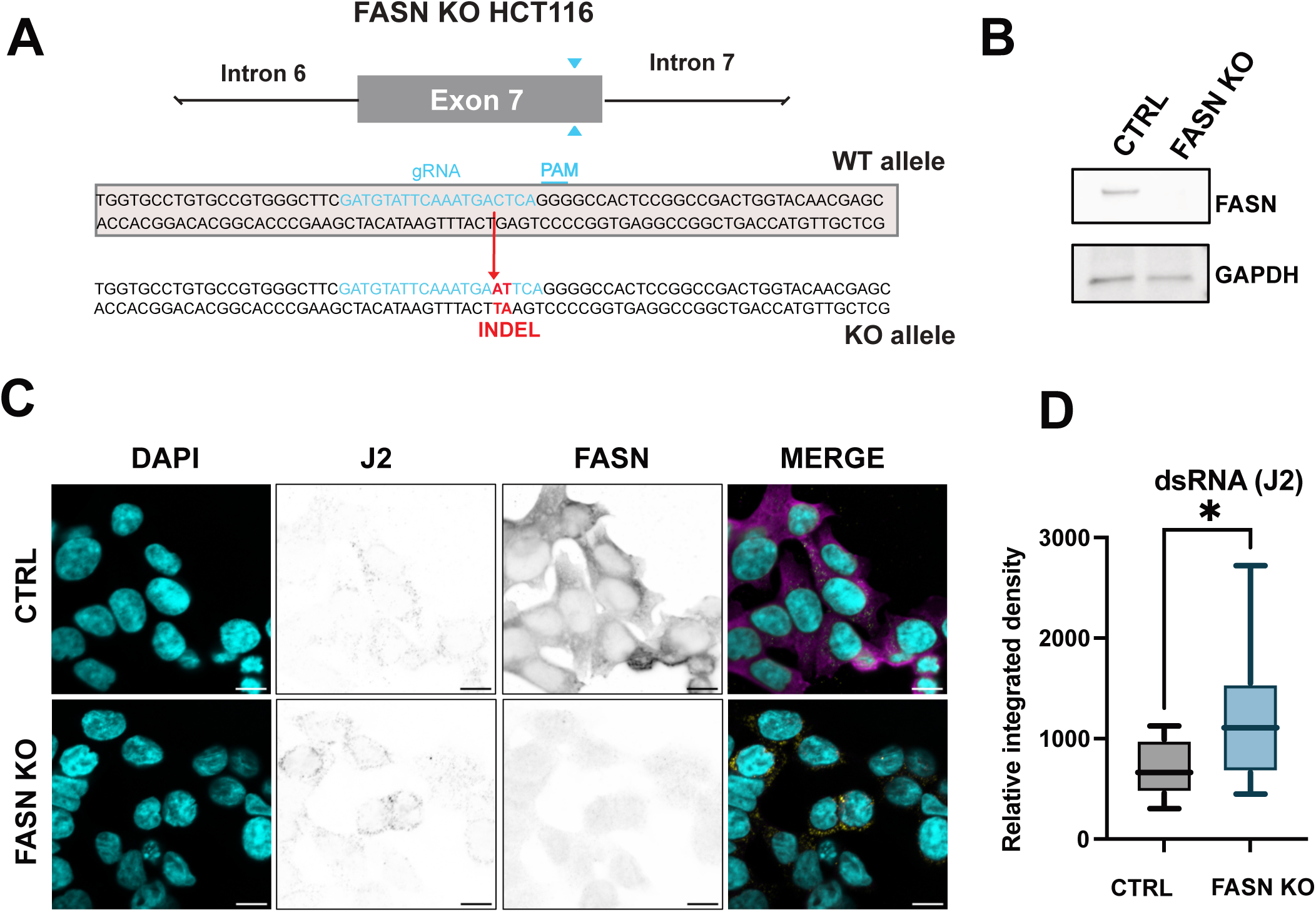
**Accumulation of endo-dsRNAs in FASN KO HCT116 cells**. **A)** Schematic representation of the FASN CRISPR/Cas9 KO (FASN KO) in HCT116 cells. One gRNA targeting the FASN exon 7 (in blue) was used to generate an INDEL which produces a substitution of G-to-A and an insertion of T (INDEL in red) causing the appearance of a premature STOP codon. **B)** Western blot on FASN and GAPDH on lysates from CTRL and FASN KO HCT116. **C)** Representative confocal co-immunostaining images using mouse J2 anti-dsRNAs (in yellow) and rabbit anti-FASN (in magenta) primary antibodies in CTRL and FASN KO HCT116 cells. DNA was stained with DAPI (cyan). Scale bar: 10µM. **D**) The relative integrated density of dsRNA staining was quantified in 20 areas per conditions using the Fiji software. Statistical analysis was performed using t-test (* = pval<0.05).

Given our prior observation that FASN might be associated with dsRNAs ^17^, we examined endogenous dsRNA levels by immunostaining using the J2 antibody ^33,34^ and observed that FASN KO HCT116 cells exhibited a significant increase in cytoplasmic dsRNA staining compared to CTRL cells (**Figure 1C-D**).

To validate these observations, we performed siRNA-mediated knock-down of FASN (siFASN) in HCT116 cells and assessed dsRNA levels by J2 immunofluorescence (**Figure S1B**). SiFASN-treated cells displayed a decrease in FASN protein expression as well as an increased cytoplasmic endo-dsRNA signal relative to siCTRL-treated cells, confirming that FASN depletion promotes dsRNA accumulation or accessibility. This phenotype was also observed to varying extent in other human cell lines, including breast cancer MDA-MB cells, hepatoma Huh7.5.1 cells and lung cancer A549 cells (**Figure S1C**). It is worth noting that cells with lower basal FASN protein expression, i.e. A549 and MDA-MB cells (**Figure S1D**), display a reduced dsRNA accumulation upon FASN depletion **(Figure S1C)**. Altogether, our data demonstrate that FASN depletion by either knock-down or knock-out strategy results in the detection of more cytoplasmic endo-dsRNAs by J2 across multiple cell lines.

### Loss of FASN rewires mitochondrial metabolism and induces inflammatory pathways in HCT116 cells

Given the accumulation of endo-dsRNA observed in the cytoplasm of FASN-depleted cells, we next investigated whether this phenotype was associated with changes in gene expression linked to immune signaling. To this end, we performed total RNA-seq on samples extracted from FASN KO and CTRL HCT116 cells. Differential gene expression analysis revealed a significant downregulation of 274 genes and upregulation of 520 genes in FASN KO cells compared to CTRL HCT116 cells (**Figure 2A and Table S1**). Gene set enrichment analysis (GSEA) revealed that downregulated genes in FASN KO cells were significantly enriched in Hallmark gene sets related to oxidative phosphorylation and MYC targets, supporting a link between FASN loss, impaired mitochondrial function and cell growth (**Figure 2B**). Consistent with these findings, gene set enrichment analysis on downregulated genes upon FASN KO shows a significant enrichment in gene ontology cellular component (GO CC) terms related to both translation and energy production **(Figure S2A)**. When looking at the gene sets enriched among the upregulated genes, we observed that apoptosis and inflammatory response, especially linked to TNF alpha signaling via NFkB, showed the highest significant enrichment scores (**Figure 2B**).

**Figure 2.**
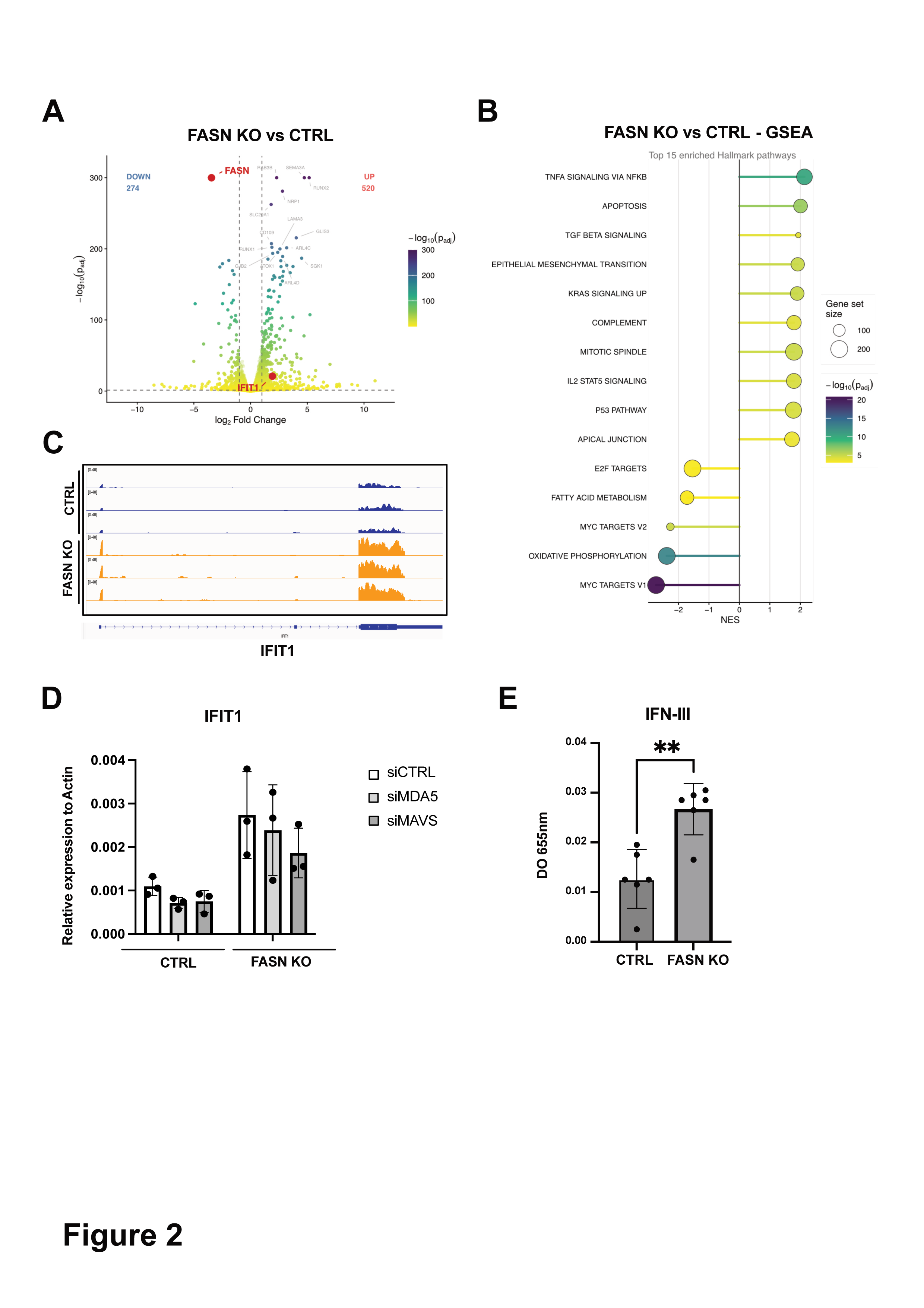
**Transcriptomic analysis of FASN KO and CTRL HCT116 cells**. **A)** Volcano plot displaying differential gene expression between CTRL and FASN KO HCT116 cells. Significantly differentially expressed genes (adjusted p-value < 0.05, |logLJFC| > 1) are colored by statistical significance (−logLJLJ adjusted p-value, blue to yellow). Vertical dashed lines indicate logLJFC thresholds of ±1. The 15 most significant genes are labeled, FASN and IFIT1 are highlighted in red. **B)** Gene set enrichment analysis (GSEA) of all expressed genes in FASN KO versus control HCT116 cells using the MSigDB Hallmark gene set collection. Genes were ranked by their DESeq2 Wald statistic. The top enriched gene-sets are displayed as enriched (positive NES, right) or depleted (negative NES, left). Color indicates statistical significance (−logLJLJ adjusted p-value, blue to yellow) and circle size represents gene set size. **C)** IGV representation of read coverage mapped to IFIT1, normalized to counts per million (CPM). Normalized read coverage is shown for CTRL (blue) and FASN KO (orange) HCT116 cells across three biological replicates. **D)** Relative IFIT1 expression in CTRL and FASN KO HCT116 cells, transfected with siCTRL, siMDA5 or siMAVS, measured by RT-qPCR. Gene expression was normalized over ACTIN housekeeping gene expression. Results represent the mean ± standard deviation (SD) of three biological replicates (n = 3). **E**) Type III IFN production in the supernatant of CTRL and FASN KO HCT116 cells monitored using HEK-Blue IFN-Lambda reporter. Reporter activity was quantified by measuring the optical density (OD) at 655 nm over three biological replicates. Statistical analysis was performed using t-test (**=pval<0.01).

Among the upregulated transcripts, IFIT1 was significantly increased in FASN KO cells compared to the control (**Figure 2B-C)**. We therefore monitored the ISG IFIT1 expression as a readout to investigate the contribution of RLR signaling. Individual KD of MDA5 and MAVS was performed and the level of IFIT1 induction was assessed (**Figure 2D and S2B - C**). First, we confirmed the induction of IFIT1 in FASN KO cells compared to CTRL cells in siCTRL-treated conditions. Although not statistically significant in our experimental setup, we also observed a reproducible negative tendency on IFIT1 expression upon MDA5 and MAVS depletion on IFIT1 sin both WT and FASN KO cells, suggesting that IFIT1 induction may depend on the MDA5–MAVS signaling axis.

To confirm that the transcriptional changes observed in FASN-depleted conditions were associated with an altered cytokine production, we measured type III IFN (IFNλ) production in the supernatants of KO and CTRL HCT116 cells using HEK-Blue IFN-lambda reporter cells. FASN KO cells secreted significantly higher levels of type III IFN compared to CTRL HCT116 cells (**Figure 2E**), suggesting a primed innate immune state.

Altogether, our data demonstrate that FASN deficiency leads to pronounced alterations in mitochondrial metabolic and inflammatory pathways. Upon FASN depletion, we also observed that IFIT1 is induced in an MDA5- and MAVS-dependent manner, likely due to enhanced sensing of endo-dsRNAs. Finally, these changes correlate with an increase in type III IFN production.

### FASN associates with cytoplasmic dsRNAs of nuclear origin without altering the global dsRNA landscape

FASN depletion has been reported to promote a primed-to-death state ^35^ and to alter mitochondrial morphology, which may contribute to the release of immunogenic mitochondrial dsRNAs. In line with these observations, MitoTracker labelling indicated a more diffuse and less intensely stained mitochondrial network in FASN KO compared to WT HCT116 cells, suggesting impaired mitochondrial integrity and organization (**Figure 3A**). We then performed mitochondria fractionation experiments to check for dsRNA proximity to mitochondria in our experimental settings. Western blot analysis confirmed the enrichment of mitochondrial markers such as MAVS and cytochrome C (CytoC) in the mitochondrial fraction (**Figure 3B**). Anti-J2 dot blot analysis on the total lysates (input), the nucleus/ cytosol (N/C) and mitochondrial fractions (MITO) revealed comparable dsRNA levels between WT and FASN KO cells, but showed a relatively higher enrichment of dsRNA in the mitochondrial fraction in FASN KO (**Figure 3C**). To confirm the double-stranded nature of these RNAs, purified RNAs from the total lysates and mitochondrial fractions were analysed. RNase III treatment resulted in a strong reduction of the J2 signal, validating that the detected species are *bona fide* dsRNAs (**Figure 3D**).

**Figure 3.**
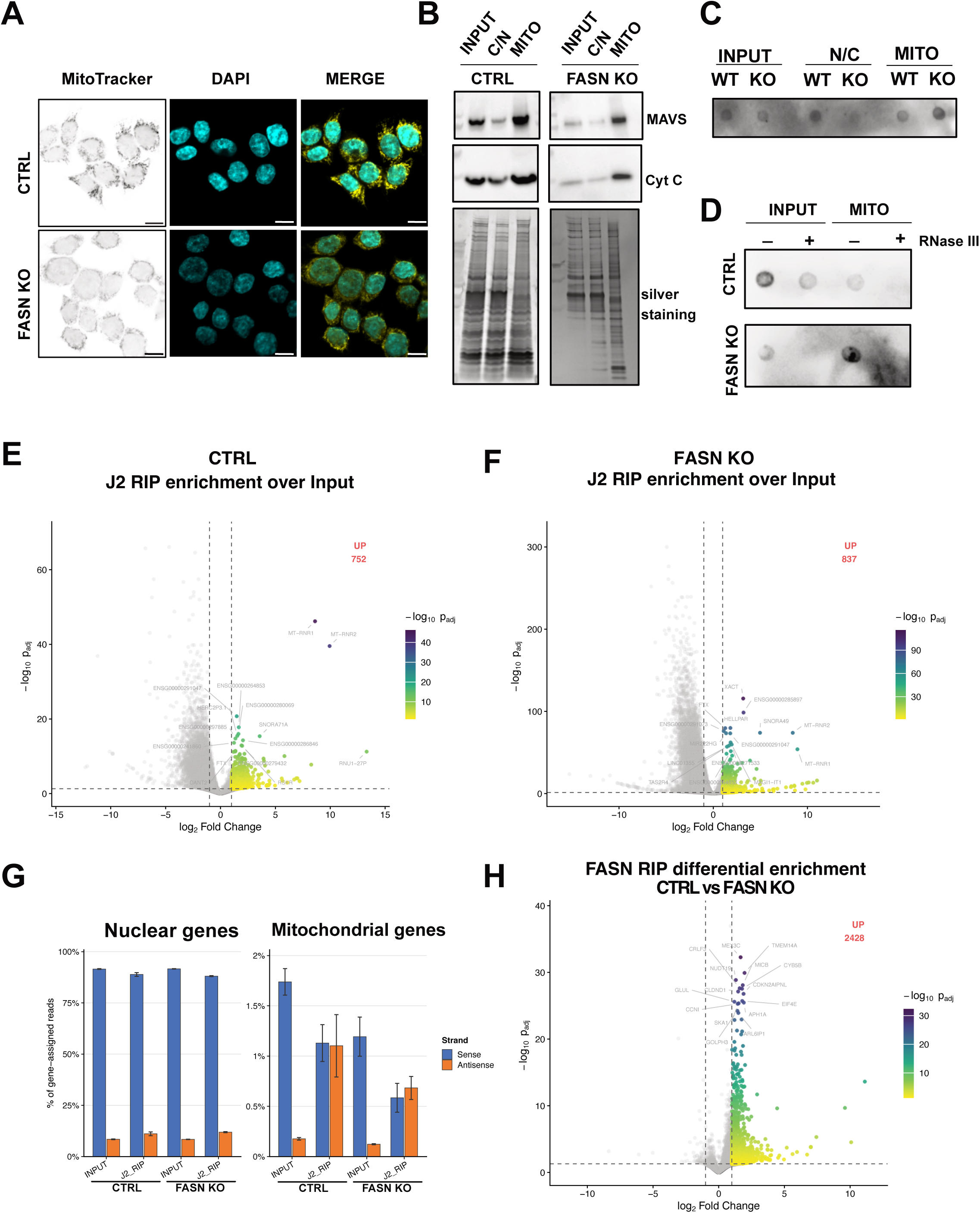
Transcriptomic analysis of FASN-associated and J2-bound dsRNAs. **A)** Representative confocal images of mitochondria in CTRL and FASN KO HCT116 cells stained using MitoTracker (yellow). DNA was stained with DAPI (cyan). Scale bar: 10µM. **B)** Western blot on MAVS and cytochrome C (Cyt C) of total lysate (INPUT), cytosol/nucleus (C/N) and mitochondrial (MITO) fractions. Silver staining is used as loading control. **C**) Anti-J2 dot blot analysis of dsRNAs on total lysates (input), nucleus/ cytosol (N/C) and mitochondrial (MITO) fractions. **D)** Anti-J2 dot blot analysis of dsRNAs from INPUT or mitochondrial (MITO) fractions in CTRL and FASN KO HCT116 cells. Purified RNA samples were treated with E. coli RNase III (RNase III +) or mock-treated (RNase III -). **E-F)** Volcano plots displaying RNA enrichment over INPUT upon dsRNA-IP-seq (J2 RIP) analysis in (**E**) CTRL and (**F**) FASN KO HCT116 cells. Significantly enriched genes (adjusted p-value < 0.05, |logLJFC| > 1) are colored by statistical significance (−logLJLJ adjusted p-value, blue to yellow). Vertical dashed lines indicate logLJFC thresholds of ±1. The 15 most significant genes are labeled. **G)** Histograms showing the percentage of reads assigned to sense (in blue) or antisense (in orange) strand of GENCODE (v47) annotated genes, split by genomic compartment (nuclear or mitochondrial genomes). Left panel: sense and antisense strand reads expressed as a percentage of all assigned reads in each sample. Right panel: sense and antisense strand reads expressed as a percentage of all assigned reads in each sample. Bars represent the mean of n = 3 biological replicates; error bars show the standard deviation. **H)** Volcano plot displaying differential RNA enrichment upon FASN-RIP-seq analysis (FASN RIP) in CTRL versus FASN KO HCT116 cells. Significantly differentially enriched genes (over input in each condition) (adjusted p-value < 0.05, logLJFC > 1) are colored by statistical significance (−logLJLJ adjusted p-value, blue to yellow). Vertical dashed lines indicate logLJFC thresholds of ±1. The 15 most significant genes are labeled.

To better characterize the nature and expression of dsRNAs and to determine whether mitochondrial RNAs could account for the dsRNA signal observed upon FASN depletion, we performed dsRNA immunoprecipitation using the J2 antibody followed by RNA sequencing (J2-RIP-seq) in control (**Figure 3E, Table S2**) and FASN KO (**Figure 3F, Table S3**) cells.

The analysis identified 752 and 837 transcripts significantly enriched over input in CTRL and FASN KO cells, respectively, indicating a broadly comparable dsRNA landscape between conditions. In both J2-IP datasets, several non-coding RNAs and two mitochondrial transcripts (MT-RNR1 and MT-RNR2, encoding the mitochondrial 12S and 16S rRNAs) were consistently detected (**Figure 3E–F**). Importantly, these mitochondrial RNAs showed similar enrichment in CTRL and FASN KO cells, indicating that they were not specifically increased upon FASN depletion and therefore did not account for the differential dsRNA signal. Their detection likely reflected the presence of stable intramolecular double-stranded structures within mitochondrial rRNAs. Consistently, strand-specific analysis revealed both sense and antisense mitochondrial transcripts within the dsRNA pool in both conditions, supporting the formation of mitochondrial RNA duplexes that are present independently of FASN expression (**Figure 3G**).

The J2-RIP enrichment over input was compared between FASN KO and CTRL cells by assessing the interaction term. Differential analysis revealed a limited number of significantly enriched or depleted RNAs (43 enriched and 22 depleted) (**Figure S3A, Table S4**), indicating that the global dsRNA landscape is largely unchanged upon FASN depletion. Of note, IFIT1 was also detected among RNAs enriched in the J2-RIP fraction in WT cells relative to FASN KO, suggesting altered association of this interferon-stimulated transcript with dsRNA-containing complexes despite its increased expression in FASN-depleted conditions.

These results suggest that the increased dsRNA signal observed by immunofluorescence is unlikely driven by mitochondrial dsRNA. Mitochondrial-encoded transcripts account for only ∼1% of total reads in the J2-RIP datasets (**Figure 3G**) and are not selectively enriched in the differential analysis between conditions. This indicates that changes in the dsRNA signal more likely reflect nuclear-encoded RNA species or other structured RNA regions.

To further determine whether FASN associates with dsRNA species, we performed RNA immunoprecipitation followed by RNA sequencing (RIP-seq) using an anti-FASN antibody in CTRL and FASN KO HCT116 cells, the latter serving as a negative control (**Figure S3B, Table S5-S6**). Assuming that all the transcripts enriched in the FASN KO background represent non-specific binders, we performed differential enrichment analysis and identified a subset of 2428 cellular RNAs specifically associated with FASN in CTRL cells (**Figure 3H, Table S7**). Notably, mitochondrial MT-RNR1 and MT-RNR2 were detected in both CTRL and FASN KO immunoprecipitations, but were not retained as differential FASN-associated targets, consistent with non-specific binding. Gene Set enrichment analysis (GSEA) showed enrichment in several cellular pathways, including mTORC1 signaling, fatty acid metabolism and oxidative phosphorylation (**Figure S3C**). Among these FASN-associated transcripts, 145 were annotated as RNA coding for mitochondrial proteins according to MitoCarta ^36^ and none corresponds to mitochondria-encoded transcripts, all being nuclear-encoded genes (**Figure S3D**).

Comparison of FASN-bound RNAs with those specifically enriched in FASN KO versus CTRL J2-RIP showed no overlap (**Figure S3E**), suggesting that the dsRNAs accumulating in FASN-depleted cells are not bound by FASN in CTRL conditions. Interestingly, approximately half of the RNAs identified as enriched in the dsRNA-IP specifically in the presence of FASN were also enriched in the FASN-RIP dataset (**Figure S3F**), suggesting that FASN associates with a specific subset of dsRNA-containing transcripts in CTRL cells. The majority of these overlapping RNAs corresponded to protein-coding mRNAs of nuclear origin involved in metabolic processes, including lipid metabolism, as well as cell cycle regulation (**Table S8**). Interestingly, IFIT1 also belongs to those RNAs being enriched in both J2- and FASN RIP in CTRL cells.

In summary, our results indicate that the absence of FASN does not affect the accumulation of mitochondrial dsRNAs, suggesting that the pre-inflammatory phenotype observed upon FASN depletion is not driven by a major mitochondrial dsRNA leakage. In contrast, FASN preferentially interacts with a subset of cytoplasmic RNAs derived from nuclear-encoded transcripts, a fraction of which was enriched in the dsRNA-RIP and is linked to lipid metabolism and cell proliferation.

### FASN association with cytoplasmic dsRNAs increases under conditions of endogenous dsRNA induction

Having established that FASN binds to a subset of cytoplasmic dsRNAs under basal conditions, we next asked whether this association was enhanced when dsRNA levels were artificially increased. To induce robust accumulation of endogenous dsRNAs of nuclear origin, we combined ADAR1 depletion with DNA methyltransferase inhibitor (DNMTi) 5-AZA treatment, a strategy previously reported to trigger immunogenic endo-dsRNA accumulation ^9,10^.

We generated ADAR1 knockout (KO) HCT116 cells using CRISPR/Cas9 and confirmed loss of both ADAR1 isoforms (constitutive p110 and IFN-inducible p150) by western blot (**Figure S4A-B).** Cells were then treated with 500 nM 5-AZA for 24 hours and harvested after four days to allow 5’AZA genomic incorporation. Under these conditions, in which cell viability was significantly reduced, as previously reported ^37^ (**Figure S4C**), RNA-seq analysis revealed a shift toward increased transcript abundance in 5-AZA treated cells compared to untreated controls (**Figure S4D-E, Table S9 and S10**), possibly due to global DNA demethylation. J2 immunostaining revealed a marked accumulation of cytoplasmic dsRNAs in ADAR1 KO cells treated with 5-AZA, which was validated by RNase III treatment (**Figure S4F-G**).

By combining ADAR1 KO cells treated with 5-AZA, we achieved sufficient endo-dsRNA accumulation to perform dsRNA immunoprecipitation followed by proteomic analysis (eDRIMS) to identify the associated proteins. Protein enrichment of the J2-IP over the IgG-IP was analyzed in WT and ADAR1 KO HCT116 cells, treated or not with 5-AZA (**Figure 4A and Figure S5A-B-C**). A total of 29 proteins were significantly enriched in the J2-IP compared to IgG controls in at least one experimental condition (**Table S11**), including known RBPs involved in endo- dsRNA regulation, such as DHX9^38^ and hnRNPC ^11^. Strikingly, FASN was among the most significantly enriched proteins in both untreated WT (**Figure S5A**) and ADAR1 KO + 5-AZA cells (**Figure 4A**). Western blot analysis confirmed that FASN, along with control RBPs such as DHX9, hnRNPU, DDX5 and hnRNPC, associated with endo-dsRNAs in all conditions tested (**Figure 4B**). These results indicate that FASN association with endo-dsRNA in untreated WT conditions is maintained in 5-AZA treated ADAR1 KO cells.

**Figure 4.**
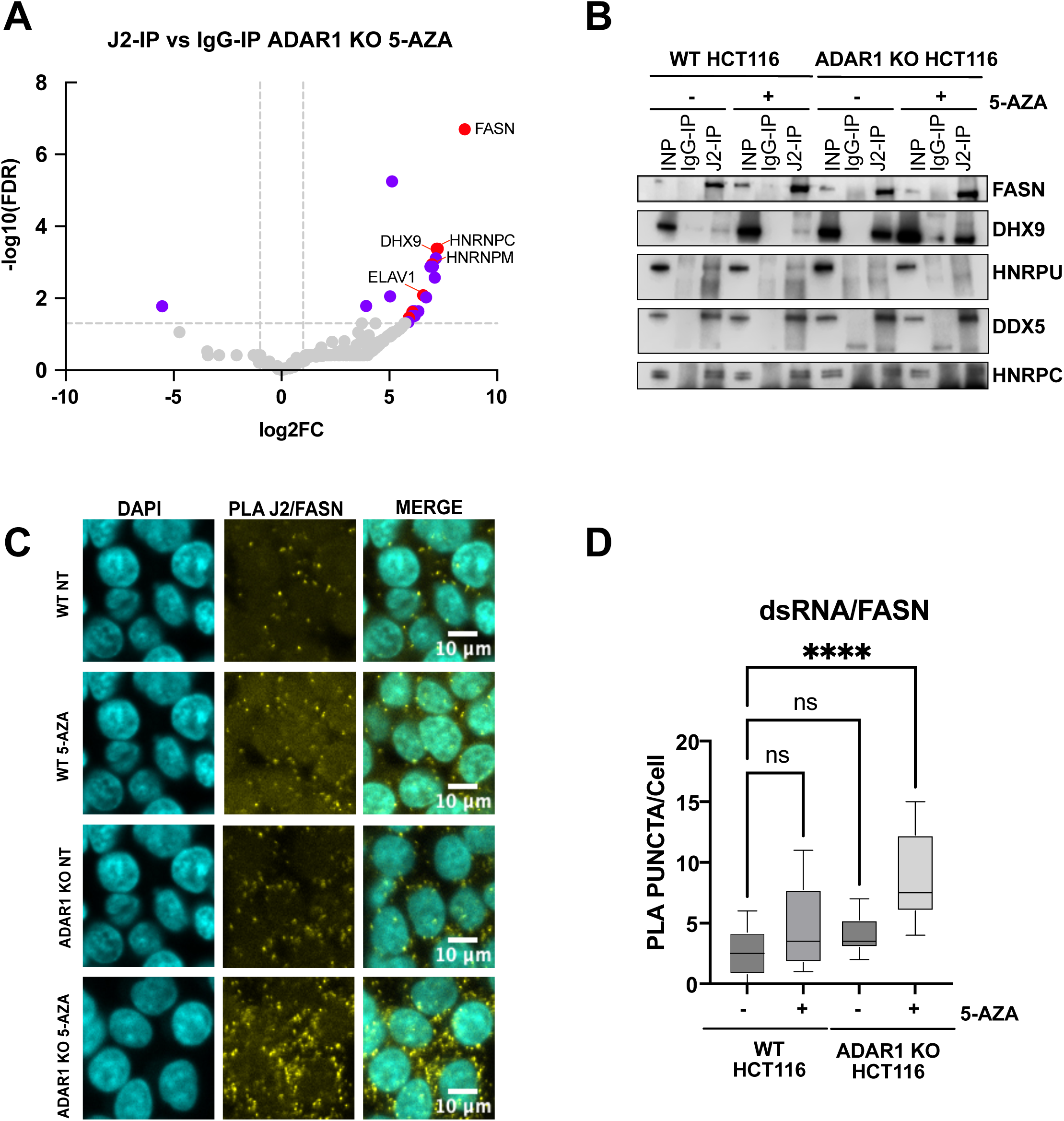
FASN association with dsRNAs increases in 5-AZA treated ADAR KO HCT116 cells. **A)** Volcano plot showing the protein enrichment identified by LC-MS/MS upon J2-IP over IgG-IP in ADAR1 KO HCT116 cells treated with 5-AZA (ADAR1 KO 5-AZA) over three biological replicates. Purple dots represent proteins that are significantly enriched (FDR>1,3, log2FC>1). Red dots correspond to proteins with a coefficient of variation (CV) <50% on the IP 5 AZA ADAR1 KO_ spectral counts. Grey dots represent non significantly enriched proteins. **B)** Western blot on pre-cleared lysates (INP), control IP (IgG-IP) or dsRNA IP (J2-IP) of WT and ADAR1 KO HCT116 cells, treated or not with 5-AZA. Primary antibodies against FASN, DHX9, hnRNPU, DDX5 and hnRNPC were used. **C)** Representative confocal microscopy images from Proximity Ligation Assay (PLA) using mouse J2 anti-dsRNAs and rabbit anti-FASN primary antibodies in WT and ADAR1 KO HCT116 cells, treated or not with 5-AZA. PLA signal is shown in yellow. DNA was stained with DAPI (in cyan), Scale bar: 10µM (n=1). **D**) Quantification of PLA puncta per cell was manually performed on 10 images for each condition. Statistical analysis was performed using one-way ANOVA test with multiple comparison to CTRL mock (**** = pval<0.0005).

Immunostaining showed that FASN is diffusely expressed in the cytoplasm in both untreated and 5-AZA-treated WT and ADAR1 KO HCT116 cells (**Figure S5D).**

To assess FASN-to-dsRNA proximity *in cellulo*, we performed proximity ligation assay (PLA) and observed that FASN was found in proximity to endo-dsRNAs even under basal conditions (up to six puncta per cell) (**Figure 4C-D**). Interestingly, PLA quantification revealed a small increase in FASN–dsRNA interactions upon 5-AZA treatment in WT cells and a larger and significant increase in ADAR1 KO + 5-AZA cells, with up to fifteen puncta per cell (**Figure 4D**). Altogether, eDRIMS and PLA results consistently demonstrate that FASN associates with nuclear-derived dsRNAs in the cytoplasm under basal conditions and that this association is enhanced under conditions of increased dsRNA accumulation, such as ADAR1 deficiency combined with 5-AZA treatment.

### FASN depletion enhances innate immune response to exogenous dsRNAs and restricts viral replication

Given the transcriptomic changes in FASN-depleted cells and the identification of FASN within the endo-dsRNA interactome, we hypothesized that FASN depletion may enhance the establishment of an antiviral-like state in response to an exogenous source of dsRNAs. To test this, we stimulated FASN KO and CTRL HCT116 cells with poly I:C and observed that FASN KO cells produced significantly higher levels of type III IFN compared to CTRL cells upon treatment (**Figure S6A**). Consistent with increased IFN production, induction of the ISGs *ISG15*, *OAS3*, *IL-18* and *IFIT1* was further enhanced in FASN KO cells relative to CTRL upon poly I:C treatment (**Figure S6B)**, indicating an increased responsiveness to dsRNA stimulation. We next assessed whether this heightened sensitivity extends to viral infection using SINV, a positive-strand RNA virus from the *Togaviridae* family, *Alphavirus* genus, which generates abundant cytoplasmic dsRNA intermediates during replication and whose viral proteins do not appear to depend on palmitoylation^39^.

Infection with a GFP- expressing SINV revealed a marked reduction in viral replication in FASN KO cells compared to CTRL, as measured by live-cell imaging (**Figure 5A**). This was corroborated by decreased viral capsid protein levels (**Figure 5B**), reduced viral genomic and sub-genomic RNA levels (**Figure 5C**) and lower viral titers (**Figure 5D).** These effects were associated with an enhanced induction of ISGs, including *ISG15*, *IL-18*, *IFIT1*, and *OAS3*, in FASN KO cells (**Figure 5E**), supporting the idea that viral restriction is due to an exacerbated antiviral response.

**Figure 5.**
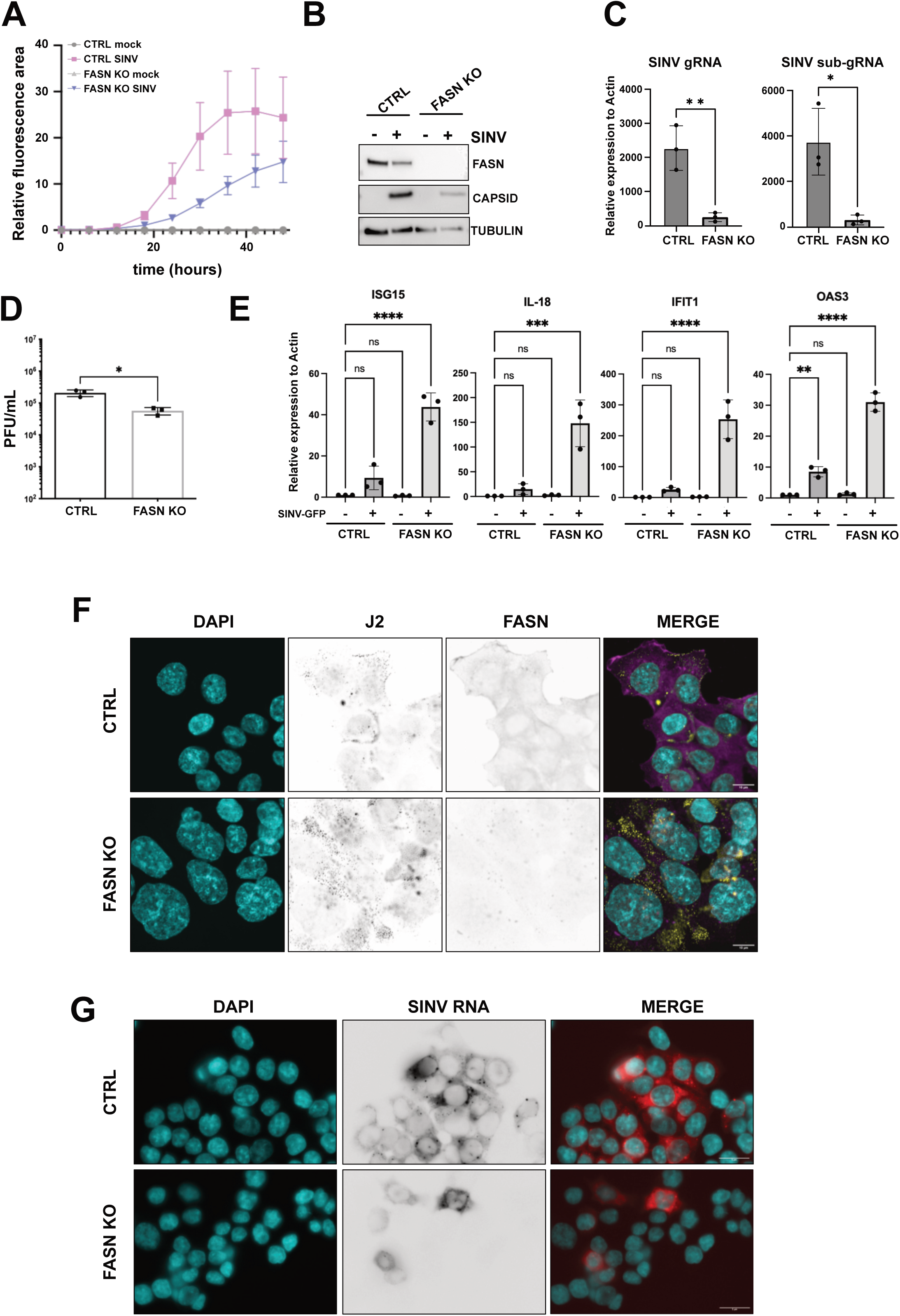
SINV infection is impaired in FASN KO cells. **A)** SINV-GFP infection kinetics in CTRL and FASN KO HCT116 cells. The relative GFP fluorescence area (expressed in percentage) as a function of time was measured after SINV GFP infection at an MOI of 0.1 every 6 h for 48 h with the CellcyteX automated cell counter and analyzer. Results represent the mean ± standard deviation (SD) of three biological replicates. **B**) Western blot on lysates from CTRL and FASN KO HCT116 infected or not with SINV-GFP MOI 0.1, 24 hours) using rabbit anti-capsid antibody. **C)** RT-qPCR on SINV genome (SINV gRNA) or sub-genome (SINV sub-gRNA) on CTRL and FASN KO HCT116 infected or not (mock) with SINV-GFP MOI 0.1 for 24 hours. Results represent the mean ± standard deviation (SD) of three biological replicates. Gene expression was normalized over actin expression. Statistical analysis was performed using a t-test (** = pval<0.01, *** = pval<0.005). **D)** Viral titers from supernatants of CTRL and FASN KO HCT116 cells infected with MOI 0.1 SINV- GFP for 24 hours as quantified by plaque assay. Results represent the mean ± standard deviation (SD) of three biological replicates. Statistical analysis was performed using a t-test (* = pval<0.05). **E)** RT-qPCR on four ISGs on CTRL and FASN KO HCT116 cells infected or not (mock) with SINV-GFP (MOI 0.1, 24 hours). Gene expression was normalized to actin expression. Results represent the mean ± standard deviation (SD) of three biological replicates. Statistical analysis was performed using a one-way ANOVA test with multiple comparison to CTRL mock (** = pval<0.01; **** = pval<0.0005). **F**) Representative confocal co-immunostaining images using mouse J2 anti-dsRNAs (in yellow) and rabbit anti-FASN (in magenta) primary antibodies in CTRL and FASN KO HCT116 cells. DNA was stained with DAPI (cyan). Scale bar: 10µM. **G**) Epifluorescence microscopy analysis of SINV (+) RNA in SINV-infected CTRL and FASN KO HCT116 cells (MOI 0.1, 24 hpi) by RNA fluorescence in situ hybridization (FISH) (in red), DAPI staining (in cyan) and merge of the two channels are shown. Scale bar: 5 μm.

Intriguingly, J2 immunostaining revealed robust dsRNA accumulation in SINV-infected FASN KO cells despite reduced viral replication (**Figure 5F**). RNA-FISH analysis of viral positive-strand RNAs confirmed a strong decrease in viral RNA levels in FASN KO cells (**Figure 5G**), suggesting that the dsRNA signal detected in these conditions is not primarily of viral origin. Instead, these observations are consistent with the accumulation of endo-dsRNAs upon infection in the absence of FASN. Altogether, our data demonstrates that FASN depletion enhances cellular responsiveness to exogenous dsRNA stimuli and restricts viral replication. The persistence of elevated dsRNA levels despite reduced viral burden further supports a model in which loss of FASN promotes the accumulation and/or accessibility of endogenous dsRNAs, thereby reinforcing an antiviral-like state.

## Discussion

FASN is the rate-limiting enzyme required for *de novo* synthesis of longLJchain saturated fatty acids (FAs) and plays a critical role in multiple cellular processes such as energy storage, membrane plasticity and protein palmitoylation ^18,25^. In addition, FASN has been characterized as a pro-viral host factor that supports viral replication through its role in lipid metabolism and the palmitoylation of viral proteins required for efficient replication ^26–29^. Independent studies support the ability of FASN to bind cellular RNAs, despite the absence of a canonical RNA-binding domain ^40,41^. Moreover, viral cross-linking and solid-phase purification (VIR-CLASP) analyses identified FASN as a pro-viral protein interacting with the viral RNA of CHIKV ^30^. These findings support the classification of FASN within a growing class of moonlighting metabolic enzymes that exhibit non-canonical RNA-binding activity ^42^. In this context, it is plausible that FASN may limit innate immune activation by sequestering immunogenic cellular RNAs away from PRRs. Conversely, association with specific RNAs could impact FASN catalytic activity and therefore influence metabolic programs linked to cellular stress responses and antiviral defense. Despite this non-canonical RNA binding capacity, the interplay between FASN, endogenous dsRNA biology and innate immune regulation remained largely unexplored.

Here, we identified FASN as a new regulator of endogenous dsRNA dynamics. We demonstrated that FASN depletion not only alters expected metabolic pathways such as fatty acid metabolism and oxidative phosphorylation, but also increases inflammation signaling. We observed that FASN depletion apparently enhances cytoplasmic accumulation of endo-dsRNA molecules but without significantly altering their global expression, indicating an effect at the level of the detection of these molecules. The observation that the dsRNA landscape is overall unchanged upon dsRNA-RIP seq between control and FASN-deficient cells supports a model in which FASN may regulate the accessibility threshold of endo-dsRNAs to PRRs rather than their expression. Although our results do not formally test the involvement of FASN metabolic activity in the observed phenotype, we hypothesize that the enhanced basal innate immune activation observed in FASN-depleted conditions is potentially driven by such immunomodulation on endo-dsRNAs.

FASN-deficient cells show enhanced production of type III interferon and increased expression of IFIT1. This response seems to depend to a certain extent on the MDA5–MAVS signaling axis, as depletion of MDA5 or MAVS attenuates IFIT1 induction in FASN-deficient cells.

At this stage, the functional importance of subcellular endo-dsRNA association in proximity to mitochondria in FASN KO cells and the link with innate immune signaling remains to be determined. observation is particularly relevant in the context of immunometabolism, in which metabolic pathways are increasingly recognized as active regulators of innate immune signaling ^43^. Mitochondria occupy a central position within this framework, as they integrate metabolic functions such as lipid metabolism and the tricarboxylic acid cycle with innate immune signaling through MAVS-dependent pathways. In this context, FASN depletion has been shown to induce a metabolic rewiring toward increased mitochondrial citrate flux and redox capacity ^44^. FASN pharmacological inhibition is also known to promote redox imbalance and increase mitochondrial apoptotic priming, reflecting a metabolically stressed state in which mitochondria become central hubs for survival and stress signaling ^35^. Consistent with this immunometabolic framework, FASN depletion in our system is associated with a primed stress state and mitochondrial-associated dsRNA localization, supporting the idea that mitochondrial rewiring may contribute to facilitate innate immune activation.

Although this spatial redistribution initially suggested a potential synthesis and leakage of mitochondrial dsRNA species in the cytoplasm, we did not observe any significant enrichment of mitochondrial-derived RNAs within the dsRNA pool in FASN-depleted conditions by RNA profiling. Our analyses further indicate that FASN preferentially associates with a population of cytoplasmic RNAs derived from nuclear-encoded transcripts, distinct from mitochondrial RNA species. A fraction of them, including IFIT1, is also enriched upon J2-RIP in the presence of FASN, suggesting that those transcripts could fold into dsRNA structures and represent candidate RNAs whose accessibility to PRRs is regulated by FASN.

These findings rather support the hypothesis that FASN directly or indirectly associates with nuclear-derived cytoplasmic RNAs and possibly regulates their accessibility to PRRs. However, so far, we have not formally demonstrated an increased engagement of endo-dsRNA by PRRs, nor their direct competition with FASN.

We showed that the degree of dsRNA detectability upon FASN depletion varies across cell types and correlates with basal FASN expression levels, supporting a dose-dependent role for FASN in limiting dsRNA exposure. Furthermore, using experimental conditions that increase endogenous dsRNA levels, such as combined 5-AZA treatment and ADAR1 depletion, we increased FASN pull-down with dsRNAs. It is important to mention that we also isolated a set of RBPs known to associate with endo-dsRNAs, such as ILF3 ^45^, HNRNPC ^11^ and DHX9 ^38^, thereby validating the robustness of pulldown approaches. Using proximity ligation assays, we observed an increased frequency of FASN–dsRNA proximity events when endo-dsRNA levels were scaled up. Together, these findings support a model in which FASN association with dsRNA increases in proportion to cellular dsRNA burden.

In our experiments, FASN depletion results in a primed antiviral state and an increased response to exogenous dsRNAs. Loss of FASN enhanced innate immunity upon poly I:C treatment and SINV, which negatively impacted replication. This antiviral phenotype is unlikely to be solely explained by defects in viral protein palmitoylation, as palmitoylation of SINV non-structural proteins is not required for replication^39^. Since dsRNA levels remain elevated despite reduced viral burden, we hypothesize that this may reflect enhanced sensing of endo-dsRNAs, ultimately leading to increased interferon responses and reduced viral replication.

The fact that FASN depletion results in an overall enhanced antiviral response is consistent with the fact that its expression is downregulated by the interferon response ^46^, suggesting a feedback mechanism linking innate immune activation to metabolic rewiring ^47–51^. Accordingly, bovine viral diarrhea virus (BVDV) infection combined with depletion of FASN enhances interferon responses, highlighting a broader connection between lipid metabolism and antiviral signaling pathways ^52,53^.

It is worth mentioning that FASN is frequently overexpressed in tumors and is a major target of anticancer therapies ^22,23^. Our results suggest that modulation of FASN activity may influence not only metabolic pathways but also endo-dsRNA biology and innate immune activation. Further characterization of the RNA species associated with FASN will be important not only to elucidate the molecular mechanisms underlying their modulatory role in immune responses in the future, but also to define its functional specificity and may open new avenues for exploiting RNA-mediated mechanisms in therapeutic strategies beyond antiviral immunity.

## Materials and methods

### Cell culture, viral stocks and virus infection

Cell lines were maintained at 37°C in a humidified atmosphere enriched with 5%CO2. HCT116 VERO E6 and HEK293T were cultured in Dulbecco’s modified Eagle medium (DMEM; Gibco; Thermo Fisher Scientific) supplemented with 10% FBS (FBS, bioSera FB1090/500), HEK-Blue IFN-Lambda cells (InvivoGen) were maintained in culture in DMEM (Gibco; Thermo Fisher Scientific) supplemented with 10% heat-inactivated FBS, 10µg/ml of blasticidin (InvivoGen), 1µg/ml of puromycin (InvivoGen) and 100µg/ml of zeocin (InvivoGen). Viral stocks available in the laboratory were produced from plasmids carrying a wild-type or a green fluorescent protein (GFP)-SINV genomic sequence, kindly provided by Dr. Carla Saleh (Pasteur Institute, Paris). Viral stocks were prepared as in ^54^. Cells were infected at an MOI of 10^-1^ and samples were harvested at 24 or 48 h post-infection as indicated in the figure legends.

### Cell treatments and transfection

For 5-AZA treatment, ADAR1 KO or WT HCT116 cells were treated with 500nM or 1µM of 5-AZA (Sigma Aldrich) or with acetic acid. After 24 h, the media was replaced by drug-free media. 72 h later, cells were harvested.

For the IFN-I treatment, ADAR1 KO or WT HCT116 cells were treated with 1000U/ml of IFN-Alpha a2 (11100-1, PBL Assay Science). Cells were harvested 24 hours later.

For poly I:C transfection, transfection complexes were prepared using poly I:C at a final concentration of either 2 or 20 µg/ml and Lipofectamine 2000 transfection reagent (Invitrogen, Fisher Scientific). The transfections were performed according to the manufacturer’s instructions and cells were harvested at 6 or 24 hours post-treatment according to figure legends.

Knock-down experiments were performed as follows: transfection complexes were prepared using siRNA targeting FASN (siFASN) or non-targeting control (siCTL) at a final concentration of 20nM and Lipofectamine 2000 transfection reagent (Invitrogen, Fisher Scientific) diluted in Opti-MEM transfection media according to the manufacturer’s instructions. The complexes were added to cells and samples were fixed for immunofluorescence 24 hours later. For MDA5 and MAVS knockdowns, transfection complexes were prepared using the specific siRNAs or non-targeting control (siCTL) (Horizon discovery, Dharmacon) at final concentration of 20nM and Lipofectamine 2000 transfection reagent (Invitrogen, Fisher Scientific) diluted in Opti-MEM transfection media according to the manufacturer’s instructions. The complexes were added to 200 000 cells in a 6-well plate and a second transfection was performed 24 hours later. Samples were recovered after 24 hours for RNA extraction.

### Cloning of sgRNA targeting FASN EX7 in pKLV

CRISPR guide RNA (sgRNA) sequences targeting the exon 7 of the human FASN gene were selected from the human sgRNA Brunello lentiviral library ^63^ sgRNA_FASN Fw: 5’CACCGATGTATTCAAATGACTCAGGT3’ sgRNA_FASN Rv: 5’TAAAACCTGAGTCATTTGAATACATC3’

Briefly, the sgRNA insert was prepared by annealing the two oligos at 95°C for 10min and then cooling down overnight. In parallel, 3µg of the pKLV-U6gRNA (BbsI)-pGKpuro2ABFP plasmid (#50946; Addgene) were digested using BbsI restriction enzyme for 4 hours at 37°C. Digested plasmid was purified on agarose gel (Monarch DNA gel extraction Kit, NEB). Diluted oligo duplexes and 100 to 300 ng of digested plasmid were ligated with the T4 DNA Fast Ligase (Thermo Fisher Scientific) using manufacturer’s instructions and DH5alpha chemo-competent *E. coli* bacteria were transformed with ligation plasmids containing an ampicillin resistance gene under sterile conditions by heat shock. Transformed bacteria were selected on LB medium + ampicillin 100µg/mL and incubated at 37°C overnight. At least one clone per conditions was cultured overnight at 37°C in 100 mL LB+ampicillin 100µg/mL. The bacteria were pelleted by centrifugation at 5000g for 15 minutes and plasmids were extracted from the bacterial pellet using midipreparations (Macherey-Nagel) and quantified by Nanodrop. Plasmid sequence was verified by Sanger sequencing (Eurofins genomics).

### Lentiviral production

Lentiviral supernatants were obtained by transfecting HEK293T cells with 1.7 µg of pKLV transfer vector carrying the indicate transgene sgRNA non targeting control or sgRNAEX7 FASN), 0.33 µg and 1.33 µg of the packaging plasmids pCMV-VSV-G (Addgene #8454) and psPAX2 (Addgene #12260), respectively, with Lipofectamine 2000 reagent (Invitrogen, Fisher Scientific). Briefly, one well from a 6-well plate with 600 000 cells was transfected for each lentiviral production. After 48 h, the medium containing viral particles was collected and filtered with a 0.45 µm PES filter and immediately used for transduction.

### Generation of CRISPR/Cas9 FASN KO HCT116 cells

Then, 100 000 Cas9-HCT116 ^64^ cells grown in blasticidine 10µg/mL were transduced in a 6-well-plate by adding either 500 µL of pKLV empty or pKLV sgRNA_FASN EX10 lentiviral supernatant to 500 µL of DMEM supplemented with 10% FBS and 4 µg/mL polybrene (Merck, Sigma-Aldrich). One day later, the medium was replaced. After 48h, 1 µg/mL puromycin was added to select the resistant cells, which were kept under constant antibiotic selection and maintained in culture with palmitic acid (P0500-10G, Sigma-Aldrich, Merck). Surviving cells were diluted in DMEM, 10%FBS to obtain 0,5 cells/well in 96-well plates and cultured for at least 3 weeks, after which genomic DNA was extracted from colonies. Cells were lysed in (50mM Tris-HCl [pH8,0]; 100mM EDTA [pH8,0]; 100mM NaCl; 1% SDS containing 0,1mg of proteinase K and incubated overnight at 55°C. Genomic DNA was extracted using phenol/chloroform/isoamyl alcohol reagent (Carl Roth) and amplified using GoTaq DNA polymerase (Promega) using specific primers surrounding the mutation site: FASN sense: 5’TGTACGCCACCTCCTGAA3’ FASN antisense: 5’TGATGCCATTCAGCTCCTGG3’ Wild-type genomic DNA was used as control template. PCR reactions were loaded in 1% agarose gel and obtained amplicons were gel purified (Monarch DNA gel extraction Kit, NEB) and sequenced by Sanger sequencing (Eurofins genomics).

### ADAR1 CRISPR/Cas9 knockout

CRISPR guide RNA (sgRNA) sequences targeting the human ADAR1 gene were taken from^53^: sgRNA#1 ADAR1 sense: 5’CACCGAAATGCTGTGCTAATTGACA3’; sgRNA#1 ADAR1 antisense: 5’AAACTGTCAATTAGCACAGCATTTC3’; sgRNA#2 ADAR1 sense:5’CACCGATGATGGCTGAAACTCACC3’; sgRNA#2 ADAR1 antisense: 5’AAACGGTGAGTTTCGAGCCATCATC3’.

Briefly, the sgRNA#1 and #2 to be inserted into the pX459V2 plasmid (#62988, Addgene) were prepared by oligos annealing at 95°C for 10 min and letting cool down overnight. In parallel, 3µg of pX45V2 were digested using BbsI restriction enzyme for 4 hours at 37°C. The digested plasmid was purified on an agarose gel (Monarch DNA gel extraction Kit, NEB). Diluted oligos duplexes and 100 ng of digested plasmid were ligated using the T4 DNA Fast Ligase (Thermo Fisher Scientific) and DH5alpha chemo-competent *E. coli* bacteria were transformed with ligation plasmids containing an ampicillin resistance gene under sterile conditions by heat shock. Transformed bacteria were selected on LB medium + ampicillin 100µg/mL and incubated at 37°C overnight. At least one clone per condition was cultured overnight at 37°C in 100 mL LB medium + ampicillin 100µg/mL. The bacteria were pelleted by centrifugation at 5000g for 15 minutes and plasmids were extracted from the bacterial pellet using a midiprep kit (Macherey-Nagel). The quantity of plasmid extracted was measured using Nanodrop. Plasmid sequence was verified by Sanger sequencing (Eurofins genomics). HCT116 cells were transfected with the two plasmids encoding for SpCas9 protein and containing sgRNA#1 and sgRNA#2, respectively. One day later, cells were treated with 1µg/ml of puromycin for 48 h. Surviving cells were diluted in DMEM, 10%FBS to obtain 0,5 cells/well in 96-well plates. Three weeks later, cellular genomic DNA was extracted from colonies. Cells were lysed in lysis buffer (50mM Tris-HCl [pH8,0]; 100mM EDTA [pH8,0]; 100mM NaCl; 1% SDS) containing 0,1mg of proteinase K and incubated overnight at 55°C. Genomic DNA was extracted using phenol/chloroform/isoamyl alcohol reagent. Then, 50ng of genomic DNA were amplified with the GoTaq DNA polymerase (Promega) using specific primers surrounding the deleted region: ADAR1 Fw: 5’GTAAGACCAGACGGTCATAGC3’; ADAR1 Rv: 5’CGCTGATGGGGTTCTTCAGC3’. Wild-type genomic DNA was used as a control template. The PCR reaction was loaded in 1% agarose gel and the obtained amplicons were gel-purified (Monarch kit, NEB) and sequenced by Sanger sequencing (Eurofins genomics).

### HEK-Blue assay

Ten thousand cells/well of HEK-blue IFN-Lambda cells (#hkb-ifnl, InvivoGen) were plated in 96-well plates in DMEM complemented with 10% heat-inactivated FBS. The day after, cells were incubated with 50µL of the supernatant of tested cell lines and quantified for 24 h. The QUANTI-Blue reagent (InvivoGen) was prepared following the manufacturer’s instruction. Next, 135µL of QUANTI-Blue were mixed to 25µL of HEK-blue IFN-Lambda supernatants in 96-well plates. The plates were incubated for 5min at 37°C and absorbance at 655nm was measured with the spectrometer iMark™ Bio-Rad.

### Protein extraction and western blotting

Cells were collected in 300 to 500µL of lysis buffer (50mM tris-HCl [pH 7,5], 150mM NaCl; 5mM EDTA, 0,05%SDS, 1%Triton X-100, 1 tablet of commercial protease inhibitor cocktail (Sigma-Aldrich) and incubated for 30min on ice. Cell lysates were collected after 15 000 g centrifugation for 5 minutes and concentration was determined using the Bradford method (Bio-Rad). After protein quantification, 30µg of protein samples were heated in 1X Laemmli buffer at 95°C for 5 minutes and separated on continuous or discontinuous (Bio-Rad), denaturing SDS-PAGE using 5% SDS-PAGE stacking gels and 10% SDS-PAGE resolving gels during 1 hour at 140V in Tris-HCl, Glycine, SDS buffer. After separation, proteins were transferred during 1 hour at 100V to 0,45µm nitrocellulose membranes (Cytiva), in Tris-HCl, Glycine, 20% Ethanol. Ponceau S staining was used to validate the protein transfer. Membranes were blocked with 5% milk-1X PBS containing 0,2% Tween 20 for 1 hour at room temperature and incubated with the indicated primary antibody overnight at 4°C. Membranes were washed three times with PBS containing 0,2% Tween 20 and incubated with indicated secondary antibody during 1 hour at room temperature. After three washes with PBS containing 0,2% Tween 20 and 1X PBS, protein signals were detected by chemiluminescence using ECL (Covalab) and Fusion FX imaging system device (Vilber).

| **Primary Antibody** | **Dilution** | **Manufacturer** |
| --- | --- | --- |
| ADAR1 | 1:1000 | CellSignaling, 812845 |
| FASN | 1:1000 | Abcam, AB128870 |
| DHX9 | 1:1000 | Proteintech, 17721-1-AP |
| HNRNPU | 1:1000 | Proteintech, 14599-1-AP |
| DDX5 | 1:1000 | Proteintech, 6702-1-Ig |
| HNRNPC | 1:1000 | Proteintech, 11760-1-AP |
| Capsid SINV | 1:2000 | Home-made, Covalab |
| GAPDH-HRP | 1:1000 | Abcam, ab2482 |
| TUBULIN-HRP | 1:1000 | Invitrogen, MA532926 |

### Mitochondrial fractionation

Mitochondria-enriched “crude” extracts were prepared as previously described in ^65^. Briefly, 10 cm2 plates were used to seed 4 million CTRL or FASN KO HCT116 cells. After 24 hours the cells were at 80% of confluency. The cells were washed twice with PBS 1X and 2 mL of MTE buffer (270mM D-mannitol, 10mM Tris base, 0.1 mM EDTA, pH adjusted to 7.4) supplemented with protease (cOmplete™ Roche, Merck) and 2µL of Ribolock (Thermo Scientific, Fisher Scientific) were added on the cells. The cells were detached using a scraper and the lysates were sonicated 3 times 10 or 30 seconds depending on the cells with the Diagenode sonicator. The lysates were centrifuged at 700g for 10 min at 4°C to remove debris. At this stage, 5 % of supernatants were kept as input samples. Then the supernatants were centrifuged again for 10 min at 15,000g and 4°C to obtain the pellets containing the “crude” mitochondria and in the upper part the nucleus and cytoplasmic fractions. The input, nucleus and cytoplasmic fractions and mitochondria-enriched fraction were used for RNAs and proteins analysis. To analyse the proteins, 12.5µL of the input (0.6%), 12.5µL (0.6%) of the nucleus and cytoplasmic fractions and 2.5µL (5%) of the “crude” mitochondria were loaded on a 4-20% acrylamide gel after adding protein loading buffer and analyzed by western blot. Cyto C (Proteintech) and MAVS antibody (Santa Cruz) were used to detect and validate the fractionation. The RNAs were extracted from 30µL of input (1.5%) and 7µL (14%) of mitochondria extracts. The RNA extraction was done with TRIzol-Reagent (Invitrogen, Fisher Scientific) according to manufacture recommendation. The RNAse treatment was done at room temperature during 30 min and 2U of RNAse III (Invitrogen Ambion, Thermo Fisher Scientific). DsRNAs were analyzed by Dot Blotting. Briefly 2µL of the different RNA samples were loaded on a Hybond NX+ membrane (GE Healthcare, Amersham) and UV crosslink (auto crosslink, Stratagene). J2 antibody was added to the membrane to detect the dsRNAs.

### Mitotracker staining and co-immunostaining

Cells were grown in 8-well LabTek slide (Merck Millipore). For mitochondria labeling, 40 000 cells were incubated with 1µM Mitotracker RedCMXRos (Invitrogen, M7512) diluted in DMEM without serum at 37°C for 30 min before fixation. Cells were fixed with 4% formaldehyde (Merck, Sigma Aldrich) diluted in PBS 1X for 15 minutes at room temperature and washed three times with PBS 1X.

For RNase III treatment, cells were incubated after fixation with PBS 1X; 0,1% Triton for 5min to permeabilize and 10U of RNase III (Invitrogen Ambion, Thermo Fisher Scientific)/well were added for 20min at room temperature. Cells were then washed 3 times with PBS 1X.

Cells were blocked using blocking buffer (0,1% Triton X-100, 5% goat serum, PBS 1X) for 1 to 3 hours at room temperature. After incubation with primary antibodies diluted in blocking buffer for 1 to 3 hours in the dark at room temperature, cells were washed three times with 0,1% Triton X-100, PBS 1X and incubated with secondary antibody coupled with Alexa-488 (A11008, Invitrogen, Fisher Scientific) and Alexa-594 (A11032, Invitrogen, Fisher Scientific) diluted at 1:1000 in blocking buffer for 1 hour in the dark at room temperature. Cells were washed 3 times with 0,1% Triton X-100, PBS 1X. DAPI (D1306, Invitrogen, Thermo Fisher Scientific) staining was performed for 5 minutes to reveal the nuclei. Slides were mounted with Fluoromount-G mounting media (Invitrogen, Thermo Fisher Scientific) and observed by confocal microscopy (LSM780, Zeiss). Images were analysed using the open-source image analysis Fiji or Image J software (1.54p) software and fluorescence intensity profiles were obtained. The mean of the relative integrated intensity in 10 to 20 cells was calculated using the same size of Region Of Interest (ROI). The images of each color channel were merged. The lookup tables (LUT) function was used to obtain the different colors. *Invert LUT command* was used to invert the colors without changing the pixel values.

| Primary Antibody | Dilution | Manufacturer |
| --- | --- | --- |
| J2 | 1:400; 1:500 | Jena BioScience |
| ADAR1 | 1:500 | Cell Signaling Technology, 812845 |
| FASN | 1:400; 1:500 | Abcam, AB128870 |
| MAVS | 1:200 | Santa Cruz, sc-166583 |

### RNA-FISH

Mock or SINV-WT infected (MOI 0.1, 24hpi) WT or FASN KO HCT116 cells were grown on 18 mm round cover glass in 12-well cell culture plates. Cells were fixed with 70% ethanol for at least 1 hour at 4°C and incubated overnight at room temperature with the SINV genome specific LGC BiosearchTechnologies’ Stellaris RNA FISH Probe diluted in RNA FISH hybridization buffer (Stellaris, Biosearch technologies). DAPI staining was performed for 30 min to reveal the nuclei (D1306, Invitrogen, Thermo Fisher Scientific). Slides were mounted on coverslips with Fluoromount- G mounting anti-fading media (Invitrogen, Thermo Fisher Scientific) and observed by epifluorescence microscopy (Olympus BX51). Images were analyzed using Image J software software 1.54P. The images of each color channel were merged. The lookup tables (LUT) function was used to obtain the different colors. *Invert LUT command* was used to invert the colors without changing the pixel values.

### Proximity Ligation Assay

*In situ* PLA was performed on fixed HCT116 cells with DuoLink PLA technology probes and reagents (Merck, Sigma-Aldrich), following the manufacturers protocol. First, cells were permeabilized with PBS + Triton X-100 0.1% for 10 min. After two PBS washes, the cells were incubated with blocking solution for 1 hour at 37°C. The primary antibodies were diluted (anti-1: 400, anti-FASN 1:400) in Duolink® Antibody Diluent according to manufacture instructions for 1 h at room temperature. The slides were washed twice for 5 min with buffer A between each step. Then, cells were incubated with the PLA PLUS and MINUS probes for 60 min at 37°C. The ligation step was performed for 30 min at 37°C. The amplification step was done during 100 min at 37°C. After two washes of 10 min with buffer B 1X, the slides were washed with wash buffer B at 0.01X for 1 minute and mounted with Duolink in situ mounting medium containing DAPI. Proximity ligation assay imaging was performed with a confocal laser scanning microscope (LSM780, Zeiss). J2-FASN PLA puncta quantification was manually performed using the FIJI software. Quantification was plotted as the number of puncta per field. Ten fields representing 10 different cells were analyzed per condition. Puncta were isolated and quantified by analyzing particle function and using the same threshold.

### MTT assay

Metabolic activity was determined by MTT assay and used to determine cellular viability. Ten thousand HCT116 WT and derived cells were cultured in 200 μl medium in a 96 well plate. Cells were incubated with 180µL of media supplemented with 20µL of the yellow tetrazole substrate compound MTT (M6494, Thermo Fisher Scientific) (5 mg/ml) for 3 hours. The reaction was stopped by adding 100 μl SDS solution (10% (w/v)) and the absorbance at 595 nm was measured after dissolution of formazan crystals using a spectrofluorometer SAFAS FLX-Xenius.

### Endogenous double-stranded RNA immunoprecipitation and proteomic analysis (eDRIMS)

Protein G Dynabeads (Invitrogen, Fisher Scientific) were resuspended in FA lysis buffer (1mM EDT1 [pH8,0], 50mM HEPES-KOH [pH7,5], 140mM NaCl, 0,1% sodium deoxycholate [w/v], 1% triton X-100 [v/v], 1 tablet of commercial protease inhibitor cocktail (Merck, Roche)). 50µL of beads per IP were blocked with 10µg/mL yeast tRNA (Invitrogen, Fisher Scientific) for 1 hour at room temperature. 10µL of uncoupled beads per IP were kept at 4°C and used for a pre-clearing of the cell lysates. Then, two micrograms of mouse anti-dsRNA J2 antibody (Jena Bioscience) or mouse anti-IgG antibody (Cell Signaling) were bound to the remaining 40µL of beads overnight at 4°C. A total of 90% confluent WT or ADAR1 KO HCT116 cells treated or not with 5-AZA in 15 cm^2^ dishes were washed with 1X cold PBS. Cells were scraped and spun at 1000 rpm at 4°C for 5 minutes. Each cell pellet was lysed in 1ml of nuclei lysis buffer (50 mM Tris-HCl [pH 8,0], 10 mM EDTA [pH 8,0], 1% SDS [w/v], 1X protease inhibitor cocktail (Merck, Roche) and incubated on ice for 10min and diluted 5-fold with FA lysis buffer containing Ribolock RNase inhibitor (Thermo Scientific, Fisher Scientific). After adding 10mM MgCl2, 5mM CaCl2, 1µL of Ribolock and 1 µL of DNase, the samples were treated for 30 minutes at 37°C. EDTA (20mM) was added to stop the reaction and a spin at 13 000 rpm at 4°C for 5min was performed. Supernatant was carefully transferred to new tubes and blocked beads for pre-clearing (see above) were added and incubated for 1 hour at 4° to pre-clear the samples. Subsequently, 40 µL of lysates out of 1mL were kept aside and treated as input samples for subsequent analyses. The remaining supernatants were added to the coupled beads (IgG or J2) and incubated at 4°C overnight. A magnetic rack was used to wash the beads. The beads were washed twice with 1mL of FA lysis buffer and twice with 1mL TE buffer (10 mM EDTA [pH 8,0], 100mM Tris-HCL [pH 8,0]). 80% of beads were kept for protein extraction by incubated it 10 min at 95°C with 40 µL 2XSDS Laemmli loading buffer (120 mM Tris-HCl [pH 6,8], 20% glycerol, 4% SDS, 0,04% bromophenol blue) and 20% of beads were kept for RNA extraction using 500 µL of TRIzo-Reagent (Invitrogen, Thermo Fisher Scientific) according to the manufacturer’s instruction. Proteins were analyzed by silver staining after separation by SDS-PAGE (Bio-Rad) using the SilverQuest Silver Staining Kit (Invitrogen, Fisher Scientific) according to the manufacturer’s advice. Protein samples were prepared for LC-MS/MS analysis as described in ^54^. Samples were analyzed with 160-minute gradients on the Qexactive + (ThermoFisher Scientific). The data were then subjected to a Mascot search with the Swissprot human bank (Mascot V.2.8, database v.2023_01) and the spectra were validated with a false positive rate (FDR) of less than 1% at PSMs and protein level, a score>25, and a validation at dataset level. The total spectral counts were then validated by statistical analyses: after a DEseq2 normalization of the data matrix, the spectral count values were submitted to a negative-binomial test using an edgeR GLM regression through R (R v3.2.5). For each identified protein, an adjusted pvalue (adjp) corrected by Benjamini–Hochberg was calculated, as well as a protein fold-change (FC). The results are presented in a Volcano plot using protein log2 fold changes and their corresponding adjusted (-log10adjp) to highlight upregulated and downregulated proteins. Due to stringent purification conditions, the number of purified proteins was low which impacted the reproducibility of the quantifications. To select candidates, on top of the significant p-values, we chose the proteins with the lowest coefficients of variation (CV<50% on the IP-KO_AZA spectral counts).

### RT-qPCR

Total RNA was extracted using Tri-Reagent (Fisher Scientific) according to the manufacturer’s instructions and quantified using the spectrofluorometer Denovix DS-11FX^+^.

For total RNA analysis, 1µg or 500ng of total RNA were treated with DNase I (Invitrogen, Fisher Scientific) and then reverse transcribed using oligo(N9) primer with the SuperScript IV reverse transcriptase (Invitrogen, Fisher Scientific) according to the manufacturer’s instructions. Quantitative Real-time (q)PCR analysis was performed and analyzed using the CFX96 touch Real-Time PCR machine (Bio-Rad). CDNA diluted 1:10 was amplified using the Maxima SYBR Green qPCR Master mix (K0253, Fisher Scientific) and specific primers at an annealing temperature of 60°C. The results were normalized to the housekeeping gene using the comparative Ct method (ΔΔCt).

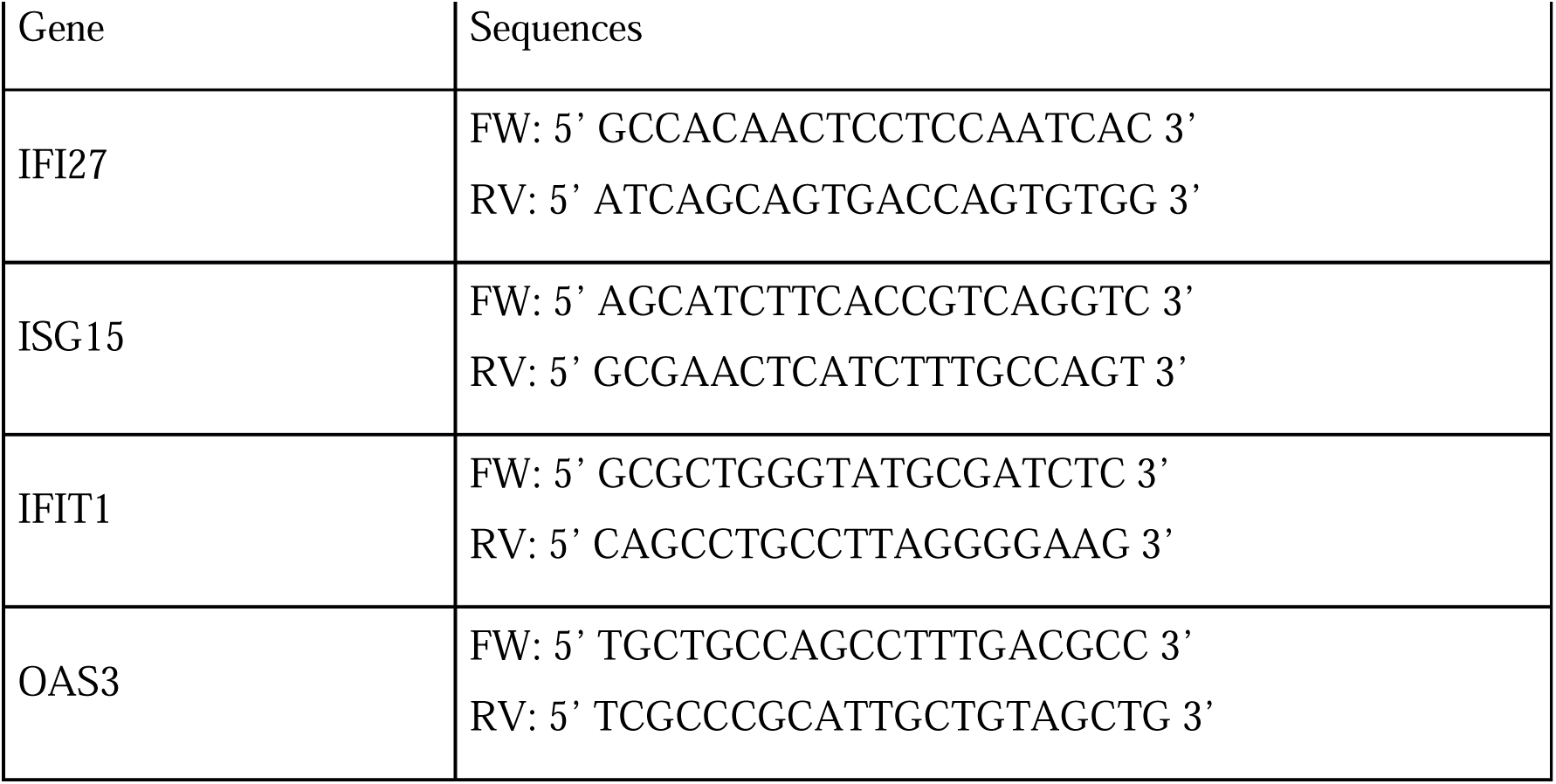
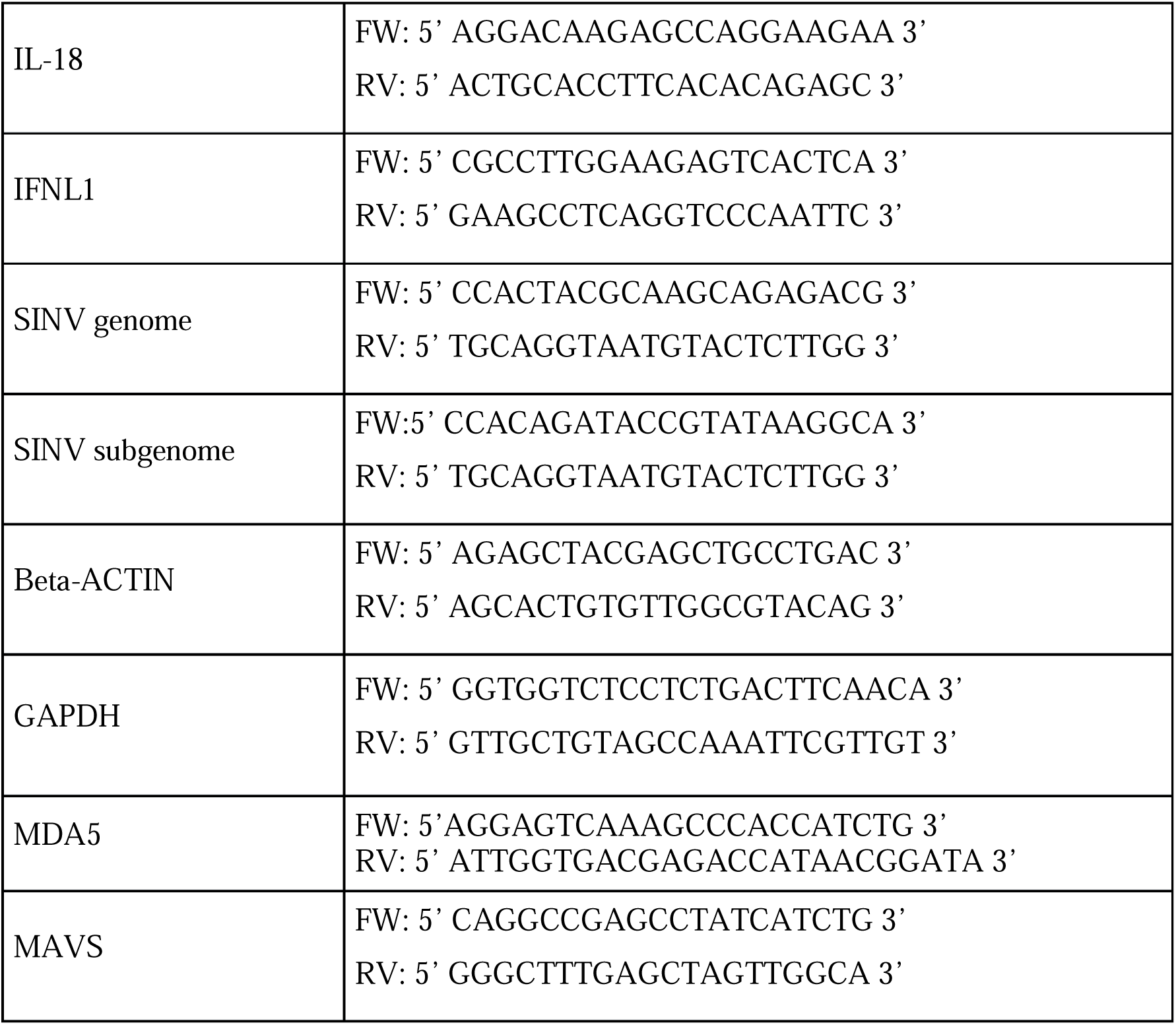

### RNA immunoprecipitation

CTRL and FASN-KO HCT116 cells were seeded in 15-cm dish to achieve approximately 80% confluence on the day of harvest. For each condition, cells from two 15-cm dishes were washed with cold PBS 1X, scraped into 1 mL PBS 1X, pooled and centrifuged at 4°C for 5 min at 1000 rpm. Cell pellets were resuspended in nuclei lysis buffer supplemented with protease inhibitors, followed by incubation on ice for 10 min. Lysates were diluted with cold FA lysis buffer supplemented with protease inhibitors. Dynabeads Protein G magnetic beads were prepared by washing twice with FA lysis buffer and blocking with BSA. Beads were coupled overnight at 4°C with either anti-dsRNA J2 antibody, anti-FASN antibody, or corresponding mouse or rabbit IgG control antibodies. Lysates were supplemented with RNase inhibitor and treated with DNase I in the presence of MgCl2 and CaCl2 at 37°C for 20 min, followed by the addition of EDTA. Lysates were clarified by centrifugation at 13,000 rpm for 10 min at 4°C and pre-cleared with uncoupled Protein G magnetic beads for 1 hour at 4°C. Pre-cleared extracts were incubated with antibody-coupled magnetic beads for 3 hour at 4°C under end-over-end rotation. Beads were subsequently washed twice with FA lysis buffer and twice with TE buffer. After the final wash, beads were resuspended in Tri-Reagent (Fisher Scientific) for RNA extraction according to the manufacturer’s instructions.

### RNA-sequencing of FASN KO vs WT

RNA from WT and FASN KO HCT116 cells in an 80% confluent 6-well plate format was extracted for library preparation and sequencing in biological triplicate. Ribo-depletion and stranded library preparation were performed at BGI Genomics (Hong Kong) and paired-end sequencing (2x 100 nt) using the DNBSEQ system was carried out by BGI Genomics (Hong Kong) to a read depth of at least 50M reads per sample. Raw sequencing reads were processed using the nf-core/rnaseq pipeline (v3.19.0) ^55^ with default parameters. Reads were aligned to the human reference genome (GRCh38, GENCODE release 47) using STAR, and gene-level counts were quantified with Salmon. Differential expression and enrichment analyses were performed with DESeq2 (v1.48.2) ^56^. Data analysis was performed on R (4.5.2) and figures using ggplot2 package (4.0.3). Genes with raw counts below 10 or TPM below 0.1 in all replicates of every experimental group were excluded prior to analysis. For the transcriptomic comparison between FASN KO and WT cells, only the input samples were used.

RNA from three independent biological replicates of J2 RIP and FASN RIP samples, from WT and FASN-KO HCT116 cells, was used for stranded RIP-RNA library preparation at BGI Genomics (Hong Kong), followed by paired-end sequencing (2x 100 nt) using the DNBSEQ system were carried out by BGI Genomics (Hong Kong) to a read depth of at least 50M reads per sample.

RIP enrichment was assessed within each condition by comparing immunoprecipitated (FASN RIP or J2 RIP) to input samples. To identify genes for which RIP enrichment significantly differed between KO and WT, a full interaction model (LJ condition + assay + condition:assay) was fitted, and the interaction coefficient was tested using the Wald test. Gene length-based normalization factors were applied across all models to account for transcript length bias. Log2 fold changes were shrunk using the apeglm method. Genes with |log2FC| > 1 and adjusted p-value < 0.05 (Benjamini-Hochberg) were considered significant. Gene Set Enrichment Analysis (GSEA) was performed using fgsea package (1.34.2), ranking all genes by their Wald statistic, against four curated collections: MSigDB Hallmark gene sets ^57^, the complete GO Cellular Component and GO Biological Process gene set collections, and Reactome pathways. To reduce pathway redundancy, a Weighted Set Cover algorithm (fgsea::collapsePathways) was applied to each collection to identify a minimal set of representative pathways. CPM-normalized strand-specific bigwig files generated by the pipeline were used for locus-level visualization of RIP and input signal tracks in the Integrative Genomics Viewer (IGV).

### RNA-sequencing of ADAR1 KO vs WT -/+ 5-AZA

RNA from WT and ADAR1 KO HCT116 cells in an 80% confluent 6-well plate format, either treated or not treated with 0.5 mM 5-AZA for 24 h, was extracted using Tri-Reagent (Fisher Scientific) according to the manufacturer’s instructions. Library preparation with the Illumina Stranded Total RNA Prep Ligation with Ribo-Zero Plus kit and paired-end sequencing (2x 100 nt) using the Illumina NextSeq 2000 system were carried out by the GenomEast sequencing platform (Strasbourg, France) to a read depth of at least 100M reads per sample on the biological triplicates. The quality and genomic origin of the sequencing reads were evaluated with FastQC (Andrews, 2010) (v0.12.1) and FastQ Screen (Wingett and Andrews, 2018) (v0.15.3). Subsequently, STAR ^58^ (v2.7.10b) was used to align the reads against the Gencode GRCh38 primary genome assembly ^59^ (v44) with the options --outSAMstrandField intronMotif, --outFilterType BySJout, and -quantMode GeneCounts. DESeq2 ^56^ (v.144.0) was used to detect differences in gene expression. Gene counts were taken directly from the STAR output and all samples were analysed together in the same object for normalisation and dispersion estimation. Only genes with a minimum average read count of 20 were included in the analysis. Differential gene expression between conditions was analysed using a two-tailed Wald test, with independent filtering disabled. The log2 fold change between conditions was estimated using Approximate Posterior Estimation ^60^. Genes with a maximum FDR of 0.01 and a minimal absolute estimated log2 fold change of 0.5 were labelled as significant. Clusterprofiler ^61^ (v4.12.0) was used for hypergeometric enrichment analysis of significant genes using the Hallmark molecular signature database ^57,62^. All genes included in the gene expression analysis were used as background genes.

### Plaque assay

To assess the amount of infectious viral particles, a total of 1 million Vero-E6 cells were seeded in a 96 well plates. The day after, the cells were infected for 1 hour with 10-fold serial dilutions of the viral supernatants. Then, the viral supernatant was removed and 100µL of a 2.5% carboxymethyl cellulose solution was added on the infected cells, which were incubated at 37°C in a humidified atmosphere of 5% CO2. Plaques were counted manually under the microscope at 48 hpi.

### Live**lJ**cell fluorescence microscopy

250 000 cells/well WT or FASN KO HCT116 cells were seeded in 12-wells plate and infected with SINV-GFP at an MOI of 0.1. Uninfected cells were used as control. GFP fluorescence and phase contrast were observed using the automated CellcyteX live-cell imaging system (Discover Echo). 4 images per well (10X objective) were acquired every 6 h for 48 h and were analyzed with the Cellcyte Studio software to determine the GFP relative intensity. The results of three biological replicates were analyzed.

### Statistical analysis

Statistical analyses were performed using GraphPad Prism 10 software. Unless otherwise stated, a paired Student’s t-test or one-way ANOVA test with multiple comparisons were performed. The number of replicates per experiment and the specific statistical tests used are stated in the figure legends.

## Data availability

The proteomics data have been deposited to the ProteomeXchange Consortium ^66^, *via* the PRIDE partner repository with the dataset identifier PXD065901.

The datasets produced in this study have been deposited in NCBI’s Gene Expression Omnibus ^67^ and are accessible through GEO Series accession number GSE307401, GSE307466 and GSE331190.

## Author contributions

**Charline Pasquier**: Conceptualization; Formal analysis; Validation; Investigation; Methodology; Writing—original draft; Writing—review and editing. **Mélanie Messmer**: Formal analysis; Validation; Investigation; Methodology; Writing—review and editing.

**Lise Moroge:** Formal analysis; Validation; Investigation; Methodology; Writing—review and editing. **Lisanne Knol**: Data curation; Bioinformatic analysis; Formal analysis; Investigation; Writing—review and editing. Lise Moroge: Formal analysis; Validation; Investigation; Writing—review and editing. **Johana Chicher**: Data curation; Bioinformatic analysis; Formal analysis; Investigation; Methodology; Writing—review and editing. **Patryk Ngondo**:

Data curation; Bioinformatic analysis; Formal analysis; Visualization; Writing—review and editing. **Sebastien Pfeffer**: Funding acquisition; Supervision; Writing—review and editing; **Erika Girardi:** Conceptualization; Supervision; Funding acquisition; Project administration; Formal analysis; Investigation; Methodology; Validation; Writing—original draft; Writing—review and editing.

## Supporting information

Genes significantly enriched in J2 RIP (contains dsRNA region) in the presence of FASN (CTRL cells) and also enriched by FASN-RIP (Associated to FASN

Differential RIP FASN enrichement over input in CTRL vs FASN KO cells (DESseq2 Interraction term analysis)

Differential RIP J2 enrichement over input in FASN KO versus CTRL cells (DESseq2 Interraction term analysis)

RIP FASN Enrichement anaysis over INPUT in CTRL cells

RIP FASN Enrichement anaysis over INPUT in FASN KO cells

RNA-seq Enrichement RIP vs INPUT samples in FASN KO cells.

RNA-seq Enrichement RIP vs INPUT samples in CTRL cells.

RNA-seq INPUT samples. Differential expression analysis on FASN KO vs CTRL cells.

RNA-seq analysis. Differential gene expression on samples ADAR1 KO HCT116 5-AZA vs NT

List of the significantly enriched proteins in the ADAR1 KO 5AZA J2 IP versus IgG IP

RNA-seq analysis. Differential gene expression on samples WT HCT116 5-AZA vs NT

## Acknowledgements

We thank all members of the laboratory for fruitful discussions, as well as Rebecca Pohen and Agathe Hunckler for technical assistance. We also thank Anne-Marie Duchene for providing Mitotracker RedCMXRos. This work of the Interdisciplinary Thematic Institute IMCbio+, as part of the ITI 2021-2028 program of the University of Strasbourg, CNRS and Inserm, was supported by IdEx Unistra (ANR-10-IDEX-0002), by SFRI-STRAT’US project (ANR-20-SFRI-0012), EUR IMCBio (IMCBio ANR-17-EURE-0023) and Equipex Insectarium ANR-11-EQPX-0022 under the framework of the French Investments for the Future Program (to SP). The mass spectrometry instrumentations were funded by the University of Strasbourg, IdEx “Equipement mi-lourd” 2015. Initial RNA sequencing was performed by the GenomEast platform, a member of the ‘France Génomique’ consortium (ANR-10-INBS-0009). CP was funded by a doctoral fellowship from the imcBio and by the Fondation ARC pour la Recherche contre le Cancer. LM was funded by a doctoral fellowship from the French Minister for Higher Education, Research and Innovation. This work received financial support from the Agence Nationale de la Recherche (EndoDRAI project ANR-22-CE15-0011-01 (to E.G.) and was supported by the Idex 2022 Attractivité grant under the framework of the IdeX University of Strasbourg (to E.G.).

## Supplementary figure legends

**Figure S1.**
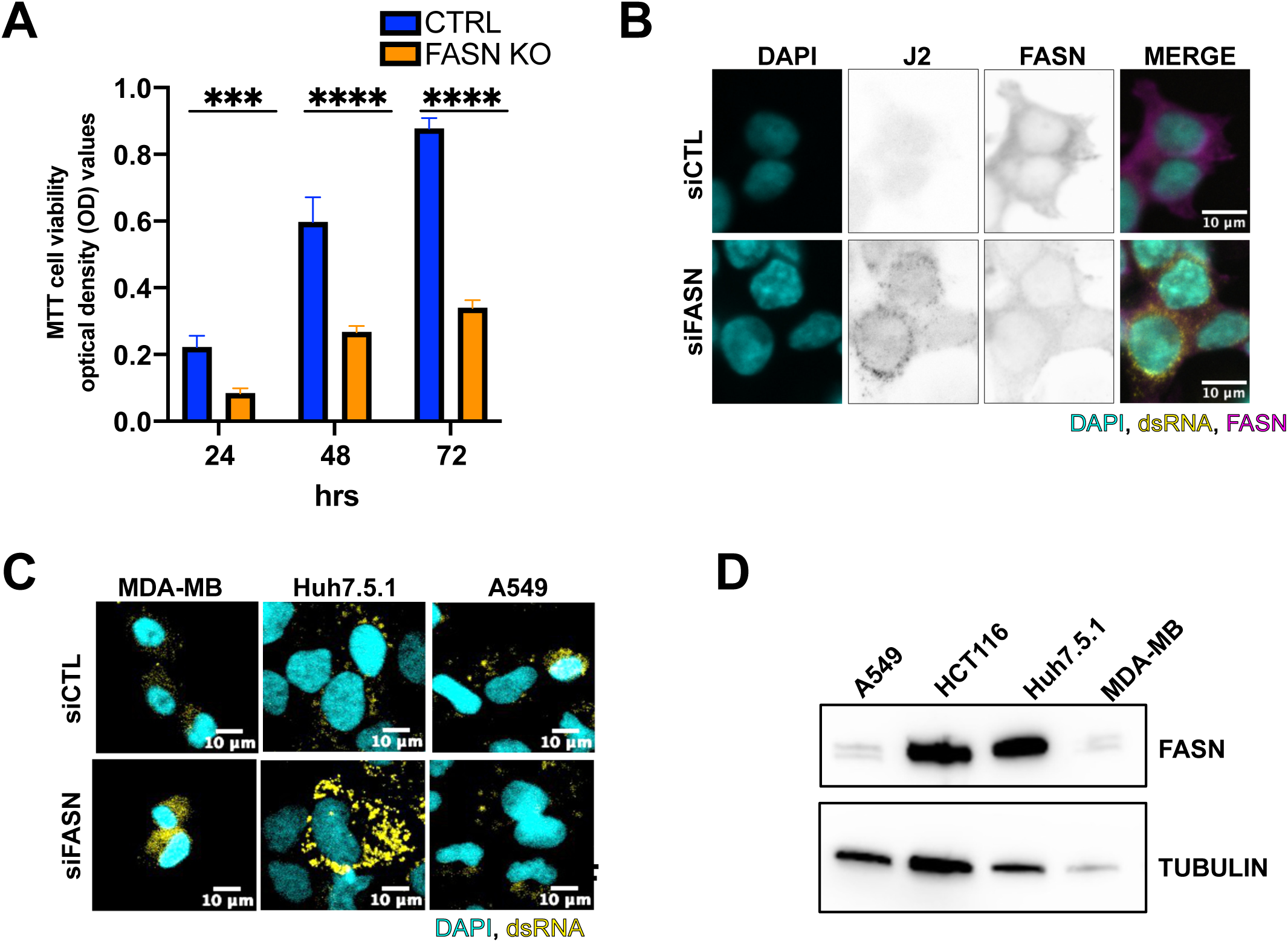
**Cell proliferation assay in FASN KO HCT116 cells and enco-dsRNA accumulation in siFASN-treated human cell lines**. **A)** CTRL (in bleu) and FASN KO (in orange) HCT116 cell proliferation monitored by MTT assay every 24 hours, over 72 hours. Optical density (OD) values were measured at an absorbance at 595 nm. Results represent the mean ± standard deviation (SD) of three biological replicates. Statistical analysis was performed using two-way ANOVA test (*** = pval<0.005; **** = pval<0.0005). **B)** Representative confocal co-immunofluorescence images from HCT116 cells transfected with anti-FASN siRNA (siFASN) or non-targeting siRNA (siCTRL) using mouse J2 anti-dsRNA antibody (in yellow) or rabbit anti-FASN (in magenta). DNA was stained with DAPI (in cyan). Scale bar: 10µM. **C)** Representative confocal immunofluorescence images from MDA-MB, Huh7.5.1 and A549 cells transfected anti-FASN siRNA (siFASN) or non-targeting siRNA (siCTRL) using mouse J2 anti-dsRNA antibody (in yellow). DNA was stained with DAPI (cyan). Scale bar: 10µM (n=1). **D)** Western blot on lysates from HCT116, MDA-MB, Huh7.5.1 and A549 cells using antibodies against FASN and TUBULIN.

**Figure S2.**
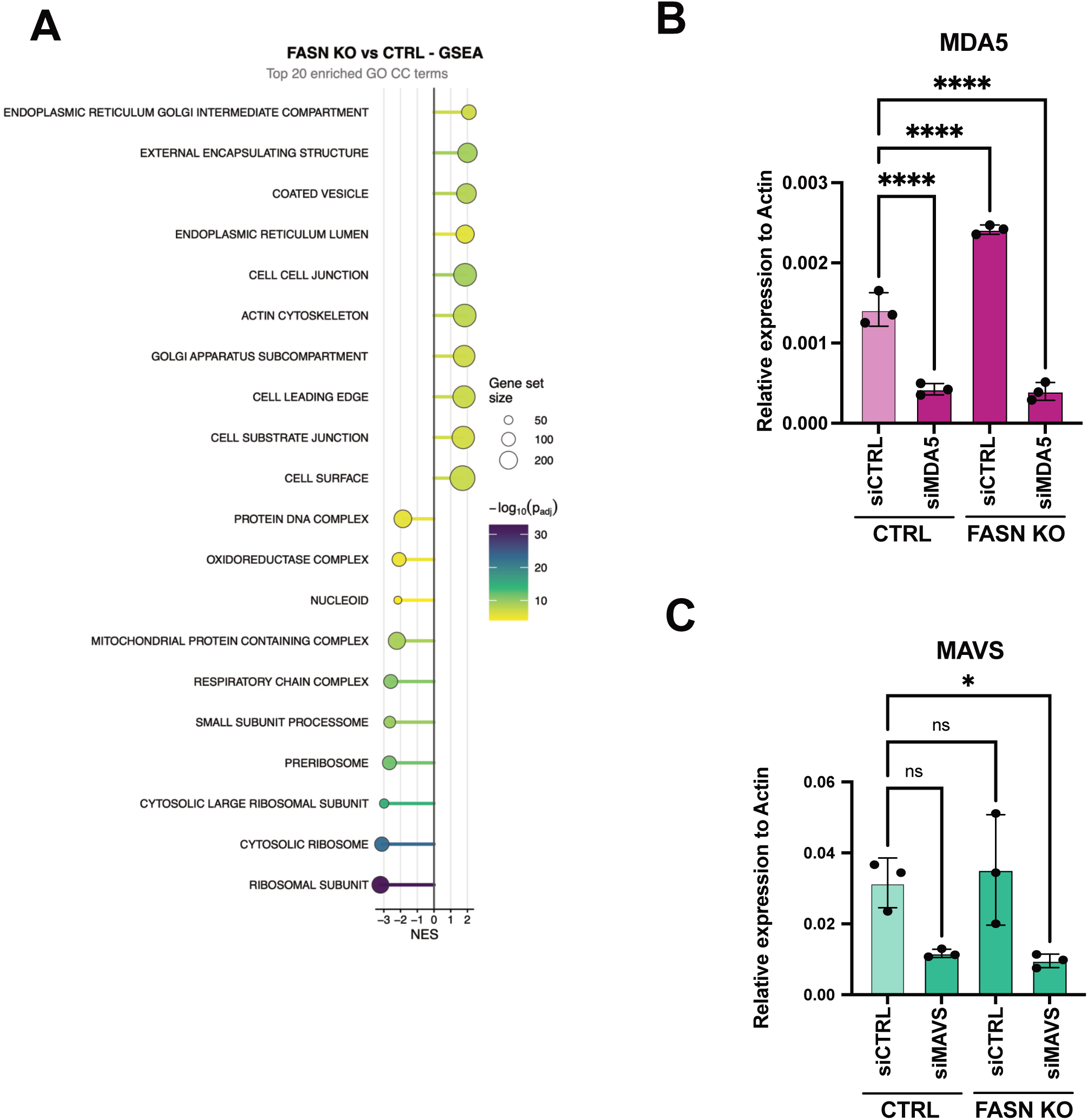
Gene set enrichment analysis of FASN KO downregulated genes using the GO cellular components and individual MDA5 and MAVS depletion in CTRL and FASN KO HCT116 cells. **A)** Gene set enrichment analysis (GSEA) of downregulated genes in FASN KO versus control HCT116 cells using the GO cellular components (CC) gene set collection. Genes were ranked by their DESeq2 Wald statistic. The top enriched gene-sets are displayed as enriched (positive NES, right) or depleted (negative NES, left). Color indicates statistical significance (−logLJLJ adjusted p-value, blue to yellow) and circle size represents gene set size. **B-C)** Relative expression in CTRL and FASN KO HCT116 cells transfected with siCTR, siMDA5 or siMAVS, measured by RT-qPCR. Gene expression was normalized over ACTIN housekeeping gene expression **B)** MDA5 and **C)** MAVS. Results represent the mean ± standard deviation (SD) of three biological replicates (n = 3). Statistical analysis was performed using two-way ANOVA test with multiple comparison to WT siCTRL (*= pval<0.05;).

**Figure S3:**
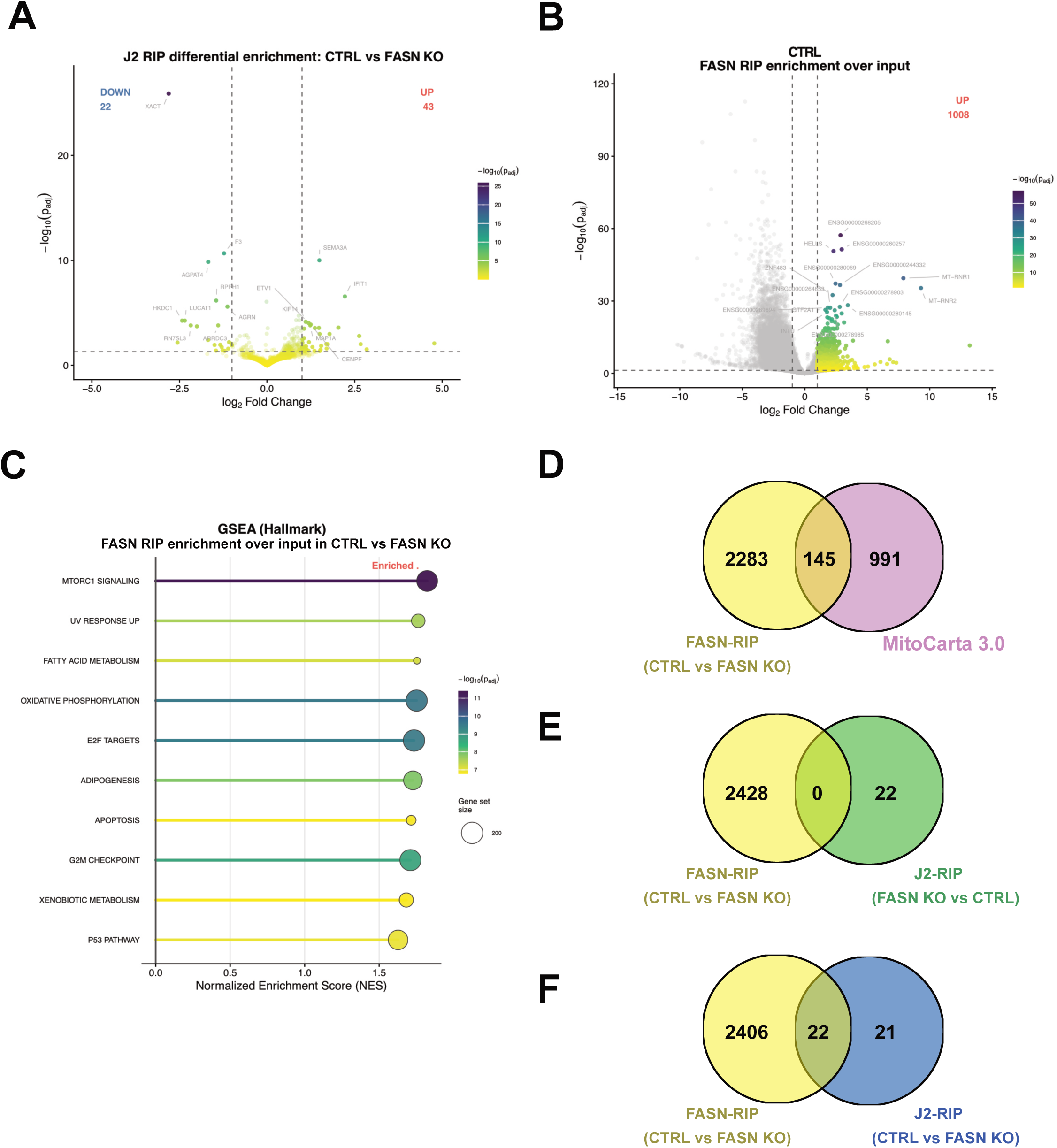
Transcriptomic analysis of FASN-associated and J2-bound dsRNA. **A)** Volcano plot displaying differential RNA enrichment upon J2-RIP-seq analysis (J2 RIP) in CTRL versus FASN KO HCT116 cells. Significantly differentially enriched genes (over input in each condition) (adjusted p-value < 0.05, logLJFC > 1) are colored by statistical significance (−logLJLJ adjusted p-value, blue to yellow). Vertical dashed lines indicate logLJFC thresholds of ±1. The 15 most significant genes are labeled. **B)** Volcano plot displaying RNA enrichment over input upon FASN-IP-seq analysis (FASN RIP) in CTRL HCT116 cells. Significantly enriched transcripts (adjusted p-value < 0.05, |logLJFC| > 1) are colored by statistical significance (−logLJLJ adjusted p-value, blue to yellow). Vertical dashed lines indicate logLJFC thresholds of ±1. The 15 most significant genes are labeled. **C)** Gene set enrichment analysis (GSEA) of significantly differentially enriched transcripts in FASN RIP (over input) in CTRL versus FASN KO HCT116 cells (related to figure 3G) using the MSigDB Hallmark gene set collection. Genes were ranked by their DESeq2 Wald statistic. The top enriched gene-sets are displayed as enriched (positive NES, right) or depleted (negative NES, left). Color indicates statistical significance (−logLJLJ adjusted p-value, blue to yellow) and circle size represents gene set size. **D-E-F)** Venn diagram showing **D**) the number of overlapping and unique significantly enriched transcripts in FASN-RIP samples (over input) in CTRL versus FASN KO (in yellow, related to Figure 3G) and genes encoding the mammalian mitochondrial proteome as annotated in MitoCarta 3.0 database (in pink), **E**) the number of overlapping and unique significantly enriched transcripts in FASN-RIP samples (over input) in CTRL versus FASN KO (in yellow, related to Figure 3G) and those significantly enriched in J2-IP (over input) in FASN KO versus CTRL (in green, related to Figure S3A, DOWN) and **F**) the number of overlapping and unique significantly enriched transcripts in FASN-RIP samples (over input) in CTRL versus FASN KO (in yellow, related to Figure 3G) and those significantly enriched t in J2-IP (over input) in CTRL versus FASN KO (in bleu, related to Figure S3A, UP).

**Figure S4:**
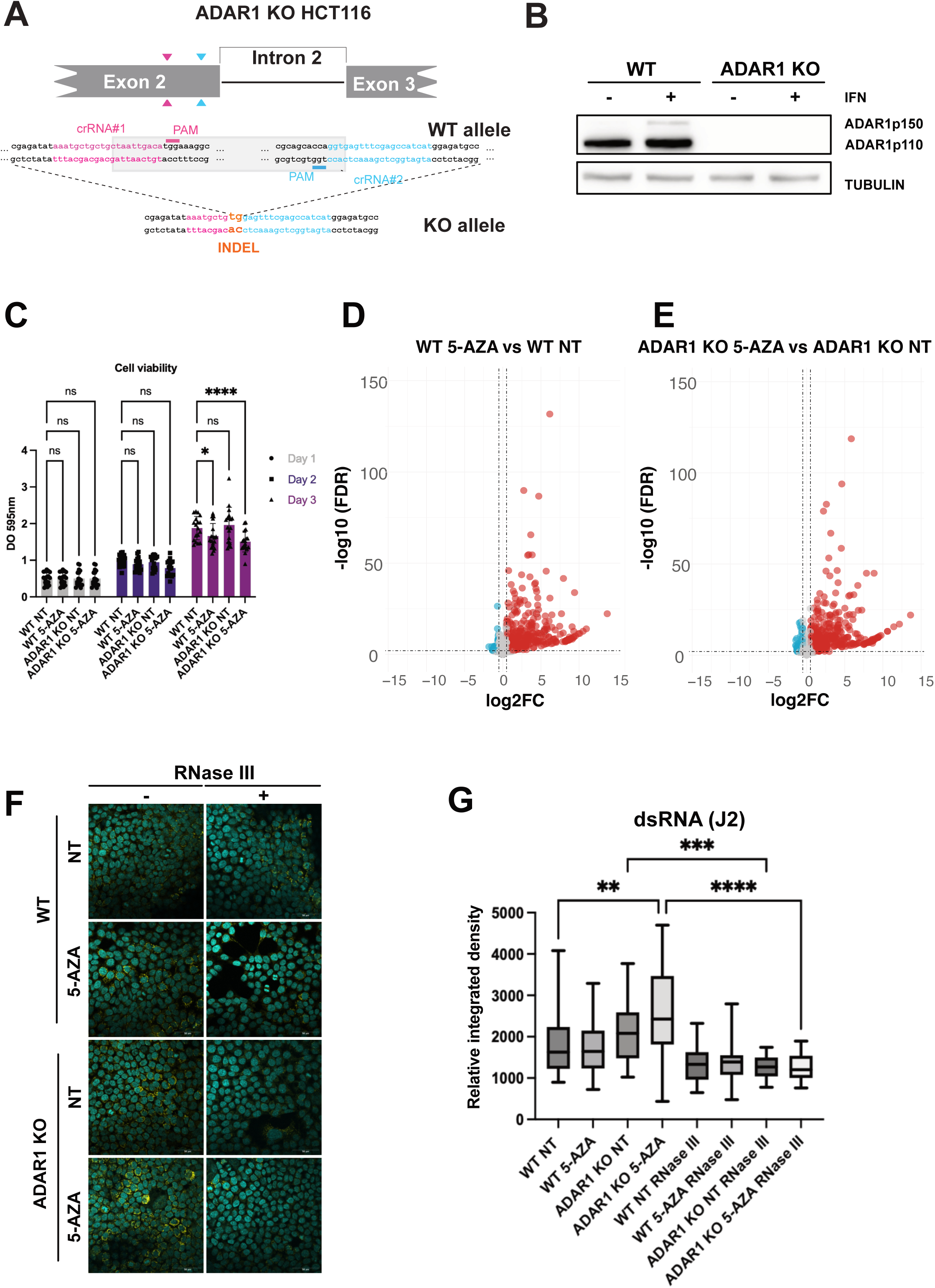
Endo-dsRNAs accumulation in ADAR1 KO HCT116 upon 5-AZA treatment with 500nM 5-AZA. **A)** Schematic representation of the ADAR1 CRISPR/Cas9 KO (ADAR1 KO) in HCT116 cells. Two sgRNAs targeting the ADAR1 exon 2 (in pink and blue, respectively) were used to delete a region of 414 bp. Indel event due to Cas9 cleavages in the KO is represented in orange. **B)** Western blot on ADAR1 and TUBULIN on lysates from ADAR1 KO and WT HCT116 with or without IFN-I treatment (1000U/mL, 24 hours treatments). **C)** ADAR1 KO and WT HCT116 cell viability monitored by MTT assay in control (NT) and 500nM 5-AZA treated (5-AZA) cells over three days. Results represent the mean ± standard deviation (SD) of three biological replicates. Statistical analysis was performed using two-way ANOVA test with multiple comparison to WT NT (*= pval<0.05; **** = pval<0.0005). **D-E**) Volcano plots displaying the differences in gene expression between either **D**) 5-AZA versus untreated WT HCT116 cells or **E**) 5-AZA versus untreated ADAR1KO HCT116 cells, over three biological replicates. Genes with an FDR value < = 0.01 and a log2 fold change >= 0.5 and are displayed in red, genes with a log2 foldchange <= -0.5 are displayed in blue. All other genes are displayed in grey. Horizontal line represents FDR = 0.01, vertical lines represent a log2 fold change of -0.5 and 0.5. **F)** Representative confocal immunofluorescence images from WT and ADAR1 KO HCT116 cells treated (5-AZA) or not (NT) with 500nM 5-AZA. Samples were treated with E. coli RNase III (RNase III +) or mock-treated (RNase III -). Staining with mouse J2 anti-dsRNA antibody (in yellow) and with DAPI (in cyan) is shown. Scale bar: 10µM. **G**) Relative integrated density was quantified in 20 areas per conditions using the Fiji software. Statistical analysis was performed using one-way ANOVA test (** = pval<0.01; *** = pval<0.005; **** = pval<0.0005)

**Figure S5.**
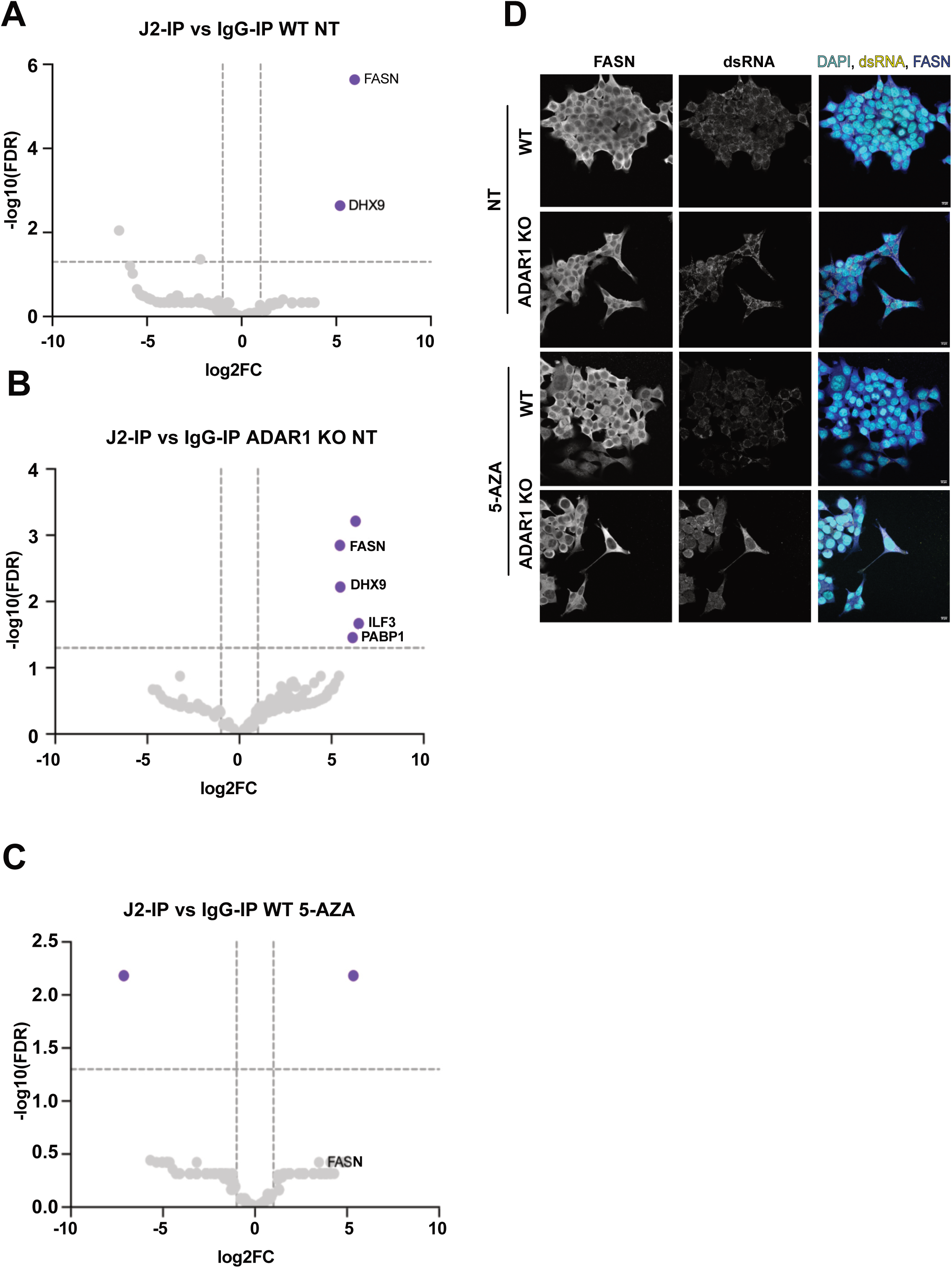
Proteome associated with endo-dsRNAs in untreated WT or ADAR1 KO HCT116 cells and 5-AZA-treated WT HCT116 cells. A-B-C) Volcano plots showing the protein enrichment upon J2-IP over IgG-IP in **A**) untreated WT HCT116 cells, **B**) untreated ADAR1 KO HCT116 cells (ADAR1 KO NT) and **C**) WT HCT116 cells treated with 5-AZA (WT 5-AZA) over three biological replicates. Purple dots represent proteins that are significantly enriched (FDR>1,3, log2FC>1). **D)** Representative confocal immunofluorescence images from WT and ADAR1 KO HCT116 cells treated (5-AZA) or not (NT) with 500nM 5-AZA. Staining with rabbit anti-FASN (in bleu), mouse anti-J2 (in yellow) and DAPI (in cyan) is shown. Scale bar: 10µM.

**Figure S6.**
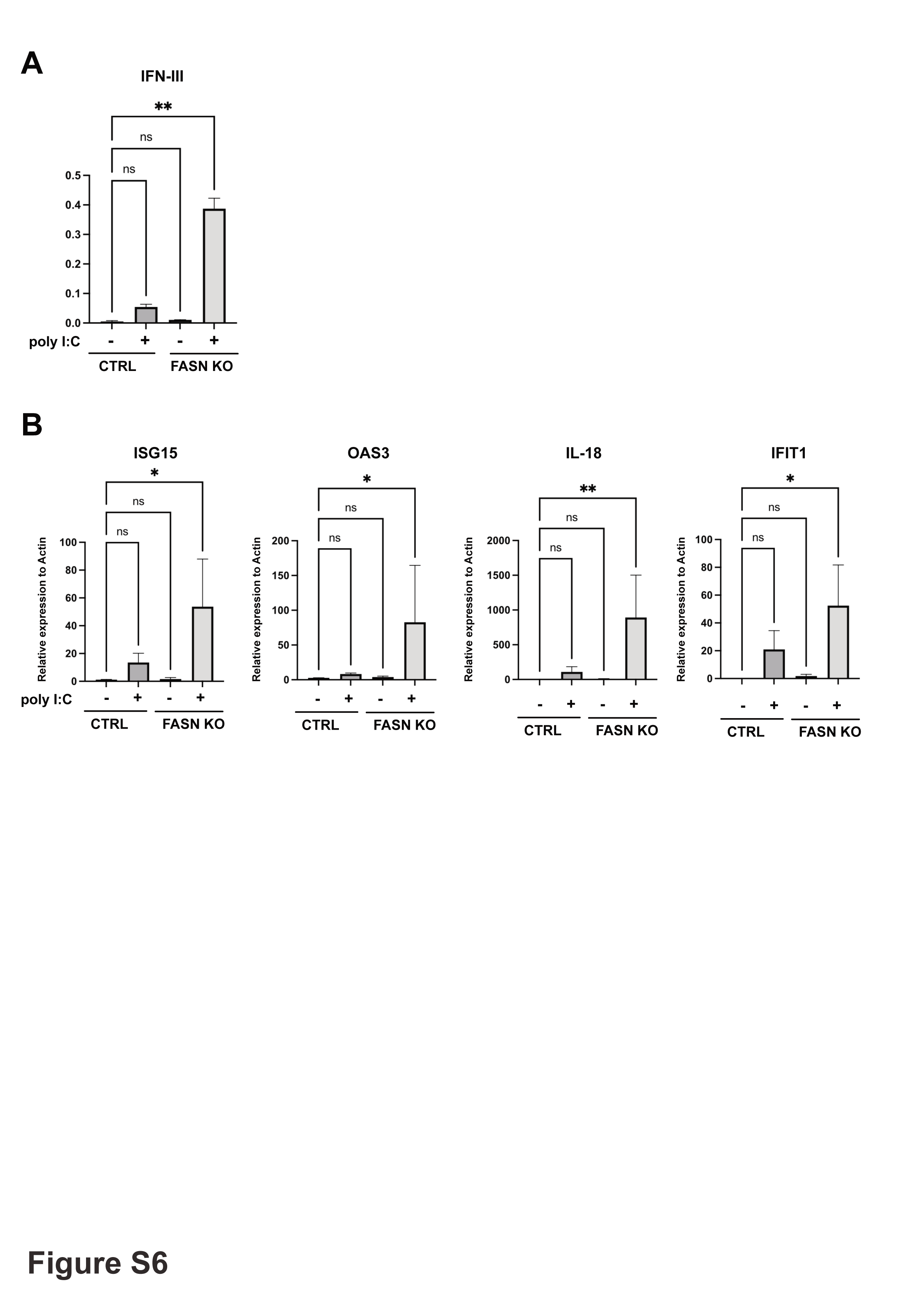
FASN KO cells show a stronger IFN response to poly I:C than CTRL HCT116 cells. **A)** Type III IFNs production by FASN KO and CTRL HCT116 cells upon poly I:C transfection (2µg/mL, 6 hours) monitored by HEK-Blue assay, OD 655nm. Statistical analysis was performed using a Kruskal-Wallis test with multiple comparison to CTRL NT (** = pval<0.01). **B)** RT-qPCR on four ISGs from FASN KO and CTRL HCT116 cells transfected with poly I:C (2µg/mL, 6 hours). Gene expression was normalized to actin expression. Results represent the mean ± standard deviation (SD) of three biological replicates. Statistical analysis was performed using a one-way ANOVA test with multiple comparison to CTRL NT (* = pval<0.05)

